# Species Context Reverses PPIP5K Control of Fungal Morphogenesis and Actin Organization

**DOI:** 10.64898/2026.08.19.745713

**Authors:** Elisa Koc, Lisa Juhran, Visnja Emmerich, Abel Alcázar-Román, Simon M. Bartsch, Adolfo Saiardi, Lasse van Wijlick, Johannes Postma, Thomas Lenz, Dorothea Fiedler, Michael Feldbrügge, Kai Stühler, Ingrid Span, Ursula Fleig

**Affiliations:** Institute for Eukaryotic Microbiology; Heinrich Heine University, Düsseldorf, Germany; Leibniz Research Institute for Molecular Pharmacology (FMP), Berlin, Germany; Medical Research Council Laboratory for Molecular Cell Biology, University College London, London, United Kingdom; Institute for Microbiology, Heinrich Heine University, Düsseldorf, Germany; Molecular Proteomics Laboratory (MPL), Heinrich Heine University, Düsseldorf, Germany; Bioinorganic Chemistry, Friedrich Alexander University Erlangen-Nürnberg, Erlangen, Germany

**Author notes:** These authors contributed equally to this work (Elisa Koc, Lisa Juhran).

## Abstract

Inositol pyrophosphates are conserved signaling molecules synthesized by the bifunctional PPIP5K enzymes, but how their cellular functions diversify across species remains poorly understood. Here, we compared the PPIP5K enzyme Asp1 in the fission yeasts *Schizosaccharomyces pombe* and *Schizosaccharomyces japonicus* and in the distantly related fungus *Ustilago maydis*. All three homologs retained a conserved kinase-phosphatase architecture and catalytic activity. However, whereas Asp1 produced broadly similar effects on actin organization and morphogenesis in *S. pombe* and *U. maydis*, its regulatory output was reversed in *S. japonicus*.

In *S. pombe* and *U. maydis*, Asp1 positively supported Arp2/3-dependent actin functions, as loss of Asp1 increased sensitivity to the Arp2/3 inhibitor CK666. In contrast, deletion of *asp1* in *S. japonicus* conferred strong CK666 resistance and caused excessive, spatially deregulated actin-patch organization. This opposing cytoskeletal phenotype was mirrored at the level of morphogenesis: Asp1 restricted the yeast-to-hypha transition in *S. japonicus*, whereas Asp1 was required for pseudohyphal growth in *S. pombe* and for filamentous development in *U. maydis*. However, the negative regulatory activity observed in *S. japonicus* was not an intrinsic property of the SjAsp1 protein. When expressed in *S. pombe*, SjAsp1 promoted invasive pseudohyphal growth, reproducing the regulatory output of the *S. pombe* Asp1 morphogenesis pathway rather than that of its native species. Similarly, SjAsp1 supported Arp2/3 functions when expressed in *S. pombe*. Thus, SjAsp1 adopted the functional behavior imposed by the host cellular environment. *S. pombe* Asp1 was originally identified as a suppressor of Arp2/3-complex mutant phenotypes, establishing a genetic connection between Asp1 and the actin nucleator. Extending this link, affinity enrichment with inositol pyrophosphates reagents recovered all seven subunits of the *S. pombe* Arp2/3 complex, providing biochemical support for a potential association between inositol pyrophosphate and Arp2/3. Together, these findings identify *S. japonicus* as a functional outlier in which a conserved PPIP5K pathway produces an opposing biological output. They further demonstrate that this divergence is determined primarily by species-specific cellular networks rather than by intrinsic differences in the Asp1 protein.

## Introduction

Inositol pyrophosphates (PP-InsPs) are highly phosphorylated signaling molecules found throughout eukaryotes, from unicellular organisms to plants and mammals. Their cellular levels are dynamically controlled by opposing kinase and phosphatase activities, allowing rapid interconversion of different PP-InsP species in response to changing cellular conditions. PP-InsPs regulate an exceptionally broad range of processes, including phosphate and energy homeostasis, transcription, stress responses, cytoskeletal organization, membrane trafficking, development and organismal physiology [1, 2]. The diversity of these functions is illustrated by their requirement for normal spermatogenesis and male fertility in mammals [3], their role in plants in linking phosphate status to beneficial mycorrhizal colonization [4], and, conversely, their manipulation by pathogenic fungi, which lower PP-InsP levels in plant cells to promote colonization [5]. These examples illustrate the remarkable biological reach of PP-InsP signaling, from reproduction and cellular homeostasis to interactions between organisms.

Despite this functional diversity, some principles of PP-InsP signaling are clearly conserved across widely separated eukaryotic lineages. Most prominently, PP-InsPs contribute to phosphate sensing and homeostasis in fungi, plants and animals, indicating that coupling PP-InsP levels to cellular phosphate status is an ancient and broadly conserved function [6–8]. In contrast, some PP-InsP-dependent functions are inherently lineage-specific, while for many others it remains unclear whether they are conserved because equivalent processes have rarely been examined comparatively.

Central to the generation of one of the major PP-InsPs, 1,5-bis-diphosphoinositol tetrakisphosphate (1,5(PP)_2_-InsP_4_), are the diphosphoinositol pentakisphosphate kinases (PPIP5Ks). PPIP5Ks are conserved bifunctional enzymes in which two opposing catalytic activities reside within one protein. Their N-terminal kinase domain phosphorylates 5-diphosphoinositol pentakisphosphate (5PP-InsP_5_) to generate 1,5(PP)_2_-InsP_4_, whereas the C-terminal phosphatase domain catalyzes the reverse reaction [9]. Structural studies of the kinase and phosphatase domains have provided insight into substrate recognition and catalysis [10–12]. By altering the balance between PP-InsP species, PPIP5Ks can modulate downstream protein functions. PP-InsPs can act through direct binding to protein targets, most prominently SPX-domain proteins [13], and have also been proposed to regulate proteins through pyrophosphorylation [14–16].

PPIP5Ks are not the only enzymes that degrade PP-InsPs. Additional phosphatases include the Siw14/PFA-DSP and Nudix families, which differ in substrate preference and phylogenetic distribution [17, 18]. Thus, cellular PP-InsP levels are determined by the balance between their synthesis and degradation.

The fission yeast *Schizosaccharomyces pombe* has become a particularly well-characterized model for PPIP5K function. Its PPIP5K family member Asp1 was originally identified through genetic interactions with the conserved actin-nucleating complex Arp2/3 [19]. Subsequent work has revealed several cellular outputs of Asp1-dependent PP-InsP signaling. Asp1-generated PP-InsPs regulate chromosome-transmission fidelity by controlling entry into mitosis, kinetochore organization, chromosome biorientation and mitotic spindle function [20–22]. Asp1 also regulates interphase microtubule organization and dynamics, with consequences for microtubule-associated mitochondrial organization [21, 23]. It further controls growth-zone selection in interphase cells and the morphogenetic switch from unicellular yeast growth to invasive pseudohyphal growth [23, 24]. Asp1-dependent PP-InsP signaling additionally regulates inorganic polyphosphate metabolism and phosphate-responsive gene expression [25–27].

Although Asp1 function has been characterized extensively in *S. pombe*, it remains unclear which of these findings represent general properties of PPIP5K signaling and which reflect relationships specific to this organism. The detailed *S. pombe* work therefore provides defined PPIP5K-dependent outputs that can be tested across species. Comparative studies of fission yeasts have already shown that conserved cellular processes can be maintained despite substantial rewiring of the underlying regulatory networks [28]. Conservation of an output in evolutionarily separated fungi would support a shared PPIP5K function, whereas divergence despite conserved PPIP5K enzymology would point to differences in how the PP-InsP signal is interpreted by the receiving cellular network. Such comparative information remains limited. Regulation of microtubule organization has been observed for PPIP5K family members in the distantly related fungi *S. pombe*, *Aspergillus nidulans* and *Ustilago maydis* [23]. For other Asp1-dependent outputs, including chromosome transmission and Arp2/3-dependent actin organization, comparable cross-species information is largely lacking. Whether PPIP5K activity promotes or restricts morphogenetic transitions in different fungi has likewise not been systematically examined. Comparative analysis can therefore distinguish conservation of PPIP5K enzymology and signal production from conservation of the cellular response.

Here, we address these questions by comparing the PPIP5K family member Asp1 in the two fission yeasts *S. pombe* and *Schizosaccharomyces japonicus* and in the distantly related basidiomycete *U. maydis*. We examine conservation of PPIP5K structure and catalytic activity and directly compare Asp1-dependent morphogenesis and cytoskeletal regulation in the three fungi. Cross-species expression distinguishes properties intrinsic to the Asp1 protein from those imposed by the cellular environment, while PP-InsP affinity enrichment coupled to quantitative proteomics investigates a potential molecular connection between 1,5(PP)_2_-InsP_4_ and the Arp2/3 actin-nucleating machinery. Together, these approaches test whether conservation of a PPIP5K signaling module extends to its biological output or whether the response to the signal is determined by the species-specific cellular network in which it acts.

## Results

### Conserved fungal PPIP5Ks retain a common domain architecture

To compare PPIP5K function across fungi, we selected the fission yeasts *S. pombe* and *S. japonicus* and the basidiomycete *U. maydis*, using the *S. pombe* Asp1 (SpAsp1) as the reference protein (Fig. 1A). *S. japonicus* diverged from *S. pombe* approximately 200 million years ago, whereas *U. maydis* belongs to the basidiomycete lineage, which separated from the ascomycete lineage approximately 600 million years ago [29–31]. PPIP5Ks contain an N-terminal kinase domain that converts 5PP-InsP_5_ to 1,5(PP)_2_-InsP_4_ and a C-terminal domain that hydrolyses 1,5(PP)_2_-InsP_4_ back to 5PP-InsP_5_ (Fig. 1B)[21, 25]. In *S. pombe*, Asp1-dependent 1,5(PP)_2_-InsP_4_ regulates a large number of biological processes including the yeast-to-pseudohyphal transition, microtubule organization and Arp2/3-dependent actin functions (Fig. 1C) [19, 23, 24].

**Fig. 1.**
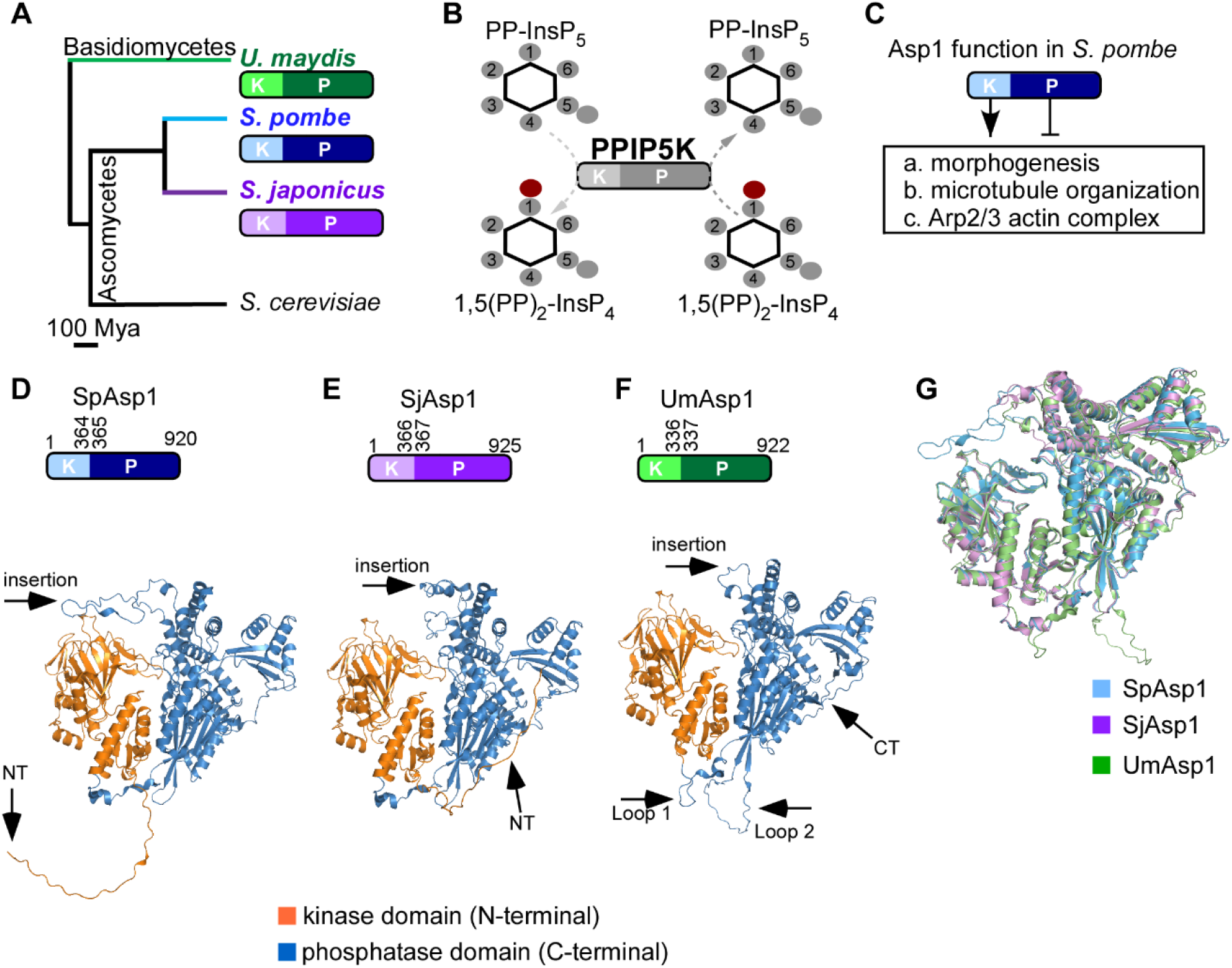
Structural comparison of the PPIP5K homologs. **(A)** Schematic representation of the phylogenetic relationships among the analysed fungal species. Tree adapted from [31]. Species-specific colours are: *S. pombe* (blue; kinase domain (K), light blue; phosphatase domain (P), dark blue), *S. japonicus* (purple; (K), light purple; (P), dark purple), and *U. maydis* (green; (K), light green; (P), dark green). **(B)** Schematic representation of the bifunctional PPIP5K enzyme. The kinase domain (K) phosphorylates 5PP-InsP_5_ to generate 1,5(PP)_2_-InsP_4_, whereas the phosphatase domain (P) hydrolyzes 1,5(PP)_2_-InsP_4_ back to 5PP-InsP_5_. Red dot represents phosphate group added or removed by Asp1 at position 1. **(C)** Overview of selected Asp1-dependent cellular processes in *S. pombe*. Positive regulation by the kinase (K) domain is indicated by an arrow, whereas antagonistic regulation by the phosphatase (P) domain is indicated by an inhibitory line **(D–F)** AlphaFold2-predicted structures of the PPIP5K family members from **(D)** *S. pombe* (SpAsp1; K domain: residues 1–364; P domain: residues 365–920), **(E)** *S. japonicus* (SjAsp1; K domain: residues 1–366; P domain: residues 367–925), and **(F)** *U. maydis* (UmAsp1; K domain: residues 1–336; P domain: residues 337–922). The predicted structures are shown as cartoon models with the N-terminal kinase domain shown in orange and the C-terminal phosphatase domain shown in blue. Black arrows indicate species-specific structural features, including the N-terminal (NT) extension in SpAsp1 (residues 1–33) and SjAsp1 (residues 1–34), the C-terminal (CT) extension in UmAsp1 (residues 877–922), an extended insertion within the phosphatase domain of SpAsp1 (residues 703–736) and SjAsp1 (residues 703– 742), which corresponds to a short loop in UmAsp1 (residues 680–683), and two UmAsp1-specific insertions: Loop 1 within the interdomain linker (residues 336–357) and Loop 2 within the phosphatase domain (residues 792–821). The confidence scores (pLDDT) and predicted aligned error (PAE) is shown in Fig.S1. **(G)** Structural alignment of SpAsp1 (blue), SjAsp1 (purple), and UmAsp1 (green) revealing similar predicted models for all organisms.

The three homologs have similar lengths: SpAsp1 comprises 920 amino acids (UniProtKB O74429), SjAsp1 (*S. japonicus* Asp1) 925 amino acids (SJAG_00244; UniProtKB B6JV42) and UmAsp1 (*U. maydis* Asp1) 922 amino acids (UMAG_06407; UniProtKB A0A0D1BUD3). Pairwise BLASTP analysis [32] showed that SjAsp1 shares 78% sequence identity with SpAsp1, whereas UmAsp1 shares 49%. AlphaFold Monomer v2.0 (AF2) models [33] revealed the characteristic PPIP5K architecture in all three proteins, with the kinase domain located at the N-terminus and the phosphatase domain at the C terminus (Fig. 1D-G).

The catalytic domains were predicted with high local confidence, as reflected by high pLDDT scores (Fig. S1). The predicted domain structures are consistent with the available structures of the human PPIP5K2 and *S. pombe* Asp1 kinase domain and the *S. cerevisiae* Vip1 phosphatase domain [10–12].

Differences among the AF2 models were concentrated in terminal extensions and low-confidence insertions within or adjacent to the predicted phosphatase domain. SpAsp1 and SjAsp1 contain short N-terminal extensions and a longer flexible insertion around residues 700 at the C-terminus, whereas UmAsp1 has a C-terminal extension and two additional loop regions, including one in the interdomain linker and one within the phosphatase domain (Fig. 1D-F; Fig. S1). These regions are predicted to be flexible and the relative orientation of the two catalytic domains is uncertain. Overall, the homologs retain a common catalytic framework but also contain species-specific flexible regions that could contribute to regulatory differences.

### SjAsp1 and UmAsp1 are required for cellular (PP)_2_-InsP_4_ production and contain conserved PPIP5K catalytic domains

To determine if the PPIP5K family members SjAsp1 and UmAsp1 were responsible for cellular (PP)_2_-InsP_4_ in these organisms, we analyzed the respective *asp1* deletion strains via SAX-HPLC analysis. Furthermore, we used a recently established PPIP5K *in vivo* screening system in *S. pombe* that allows a read-out of functional kinase and phosphatase activity [34]. Using SAX-HPLC, we determined PP-InsP levels in wild-type and *asp1Δ* strains of *S. japonicus* (yeast form) and *U. maydis* (sporidia (yeast form)). We had previously generated an *Um asp1Δ* strain [23]. For *S. japonicus*, a *Sj asp1Δ* strain was constructed in an *ura4-D3* strain by replacing the *asp1^+^* open reading frame with an *ura4^+^* cassette via homologous recombination. Next, *Sj* WT and *asp1Δ* strains and *Um* WT and *asp1Δ* strains were precultured in their respective minimal media. Cells were subsequently metabolically labeled with [³H]-inositol for approximately 5 (*S. japonicus*) or 6 (*U. maydis*) cell divisions, followed by acidic extraction of PP-InsPs and analysis by SAX-HPLC. The analysis of the *Sj* WT strain revealed three distinct peaks corresponding to InsP_6_, PP-InsP_5_, and (PP)_2_-InsP_4_ (Fig. 2A, top panel and enlargement in bottom panel). In contrast, the *Sj asp1Δ* strain lacked a detectable (PP)_2_-InsP_4_ peak and exhibited an increase in PP-InsP_5_ level (Fig. 2A, zoom). Thus, SjAsp1 is required for the production of (PP)_2_-InsP_4_ in *S. japonicus*. Next, we analyzed the *U. maydis* HPLC. The *Um* WT strain displayed a distinct InsP_6_, PP-InsP_5_, and (PP)_2_-InsP_4_ profile (Fig. 2B top panel and enlargement in bottom panel), whereas the *Um asp1Δ* strain lacked detectable (PP)_2_-InsP_4_ (Fig. 2B, zoom). Surprisingly, PP-InsP_5_ was exceptionally high in the *Um asp1Δ* strain and surpassed the InsP_6_ peak (Fig 2B, bottom panel, zoom). This is highly unusual, as InsP_6_ is typically the most abundant highly phosphorylated soluble inositol phosphate species detected by anion-exchange HPLC [35]. The SAX-HPLC data are usually normalized by performing the ratio of a specific PP-InsP over its precursor or the abundant and metabolic stable InsP_6_ [35]. Quantification of the *S. japonicus* HPLC results is shown in Fig. 2C. For *U. maydis*, the measured PP-InsP_5_ and (PP)_2_-InsP_4_ levels were also normalized to InsP₆ levels, demonstrating a severe reduction of (PP)_2_-InsP_4_ (Fig. 2D, left panel), and a significant increase in PP-InsP_5_ levels (Fig. 2D, right panel). Thus, our data demonstrate that (i) the PPIP5K homologs from *S. japonicus* and *U. maydis* are required for (PP)_2_-InsP_4_ synthesis and (ii) that the absence of UmAsp1 results in an exceptional increase of PP-InsP_5_ level not observed for any other studied eukaryote.

**Fig. 2.**
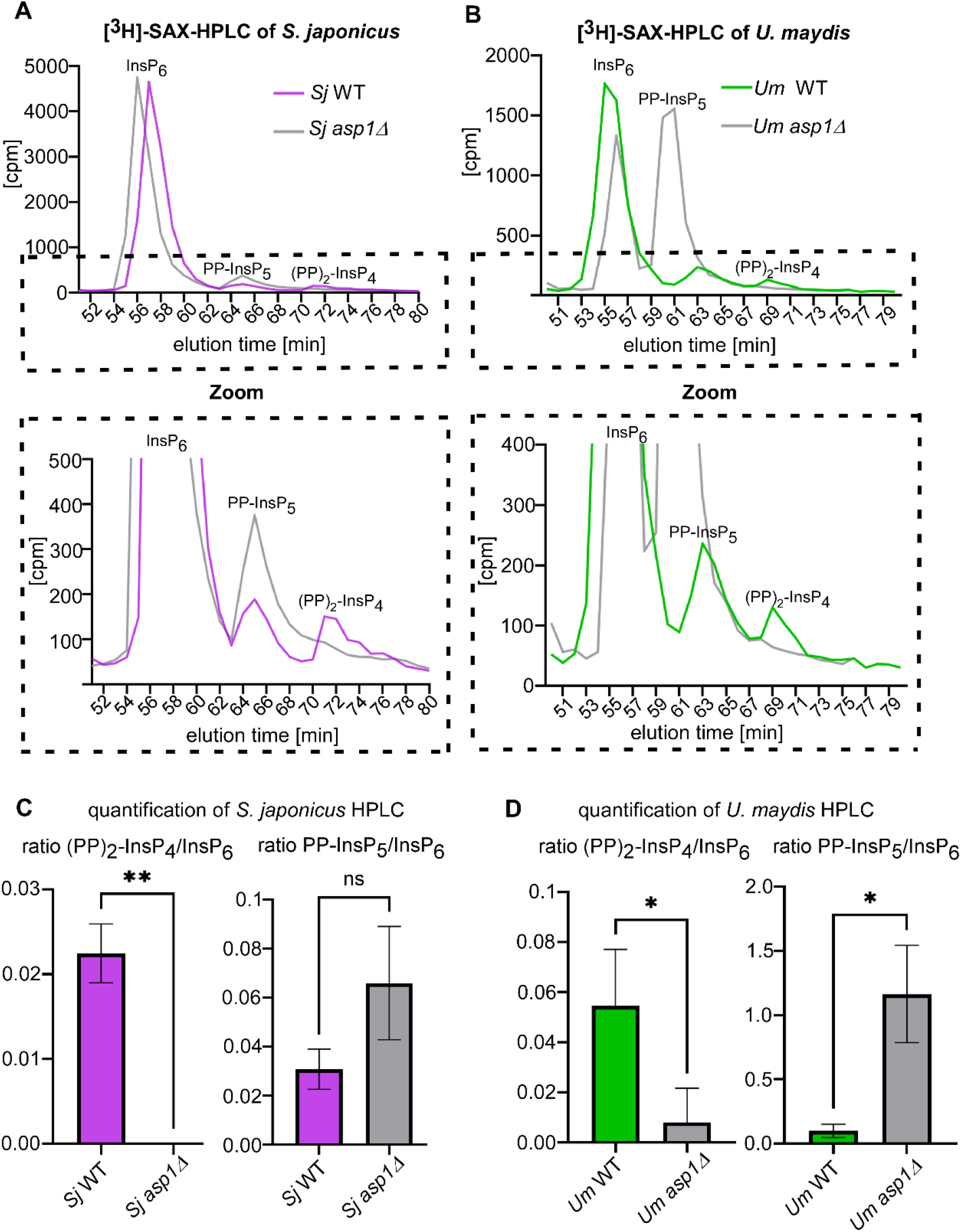
S. *japonicus* and *U. maydis* Asp1 proteins generate (PP)_2_-InsP_4_. **(A)** SAX-HPLC chromatograms of wild-type and *asp1Δ* cells from S*. japonicus* (top panel), dashed box: zoom of the PP-InsP_5_ and (PP)_2_-InsP_4_ peaks of *S. japonicus* (bottom panel) (**B**) SAX-HPLC chromatograms of wild-type and *asp1Δ* cells from *U. maydis* (top panel), dashed box: zoom of the PP-InsP_5_ and (PP)_2_-InsP_4_ peaks of *U. maydis* (bottom panel). For A and B representative SAX-HPLC examples are shown. (**C**) Quantitative analysis of (PP)_2_-InsP_4_ /InsP_6_ ratio (left panel) and PP-InsP_5_/InsP_6_ (right panel) ratio of *S. japonicus* strains. (**D**) Quantitative analysis of (PP)_2_-InsP_4_/InsP_6_ ratio (left panel) and PP-InsP_5_/InsP_6_ ratio (right panel) of *U. maydis* strains. (**C+D**) Statistical comparisons were performed using Welch’s t-test based on three independent biological replicates per strain and organism. ns, not significant; *, *P* < 0.05; **, *P <* 0.01. cpm= counts per minute.

Next, we investigated the kinase and phosphatase activities (Fig. 3A) of SjAsp1 and UmAsp1 using a recently established *in vivo* assay in *S. pombe* [34]. Briefly, elevated 1,5(PP)_2_-InsP_4_ levels confer resistance to the microtubule-destabilizing drug thiabendazole (TBZ), whereas reduced 1,5(PP)_2_-InsP_4_ levels increase TBZ sensitivity in a *S. pombe* strain [20, 21, 23, 24, 34]. Accordingly, expression of a PPIP5K kinase domain increases TBZ resistance, while expression of a PPIP5K phosphatase domain has the opposite effect (Fig. 3B-C). Plasmids encoding the kinase domains of Asp1 from *S. pombe* and *S. japonicus* already existed [34], while a codon-optimized variant from *U. maydis* was generated. All Asp1 variants were expressed on plasmids via the repressible *S. pombe nmt1^+^* promoter [36]. To assess kinase activity, wild-type *S. pombe* cells were transformed with either the control vector or plasmids expressing the respective kinase variants. Expression of all three different Asp1 kinase variants when expressed under high-expression conditions rescued the growth on TBZ-containing media (compared to control plasmid) (Fig. 3D). Thus, we conclude that they have *in vivo* kinase activity [34]. Next, we analyzed the function of the different phosphatase domains. Plasmids encoding the *S. pombe* and *S. japonicus* phosphatases already existed [34] and a plasmid encoding codon-optimized *U. maydis* phosphatase was generated. We had found previously, that analysis of PPIP5K phosphatases from other organisms than *S. pombe*, were best studied in the *S. pombe asp1^H397A^* strain which has higher than wild-type 1,5(PP)_2_-InsP_4_, due to a non-functional phosphatase domain of the endogenously encoded protein [21, 34]. *Asp1^H397A^* cells were transformed with either the control vector or plasmids expressing the respective Asp1 phosphatase domains via the *nmt1^+^* promoter. Transformants expressing SpAsp1 or SjAsp1 phosphatase showed reduced growth on TBZ-containing medium, demonstrating functional phosphatases (Fig. 3E, top panels). However, expression of the UmAsp1 phosphatase under these high-level expression conditions, resulted in a non-growth phenotype of the transformants even on media without TBZ (Fig. 3E, top panels, high expression). Thus, to be able to assess UmAsp1 phosphatase activity, we grew transformants on media, which did not allow full de-repression of the *nmt1^+^* promoter. Under these conditions, growth of transformants expressing UmAsp1 phosphatase was not affected on control media (Fig. 3E, lowest left panel), but growth was reduced on TBZ-containing plates (Fig. 3E, lowest right panel). We conclude that in our *S. pombe in vivo* assay, the Asp1 kinase and phosphatase domains from all three organisms show activity.

**Fig. 3.**
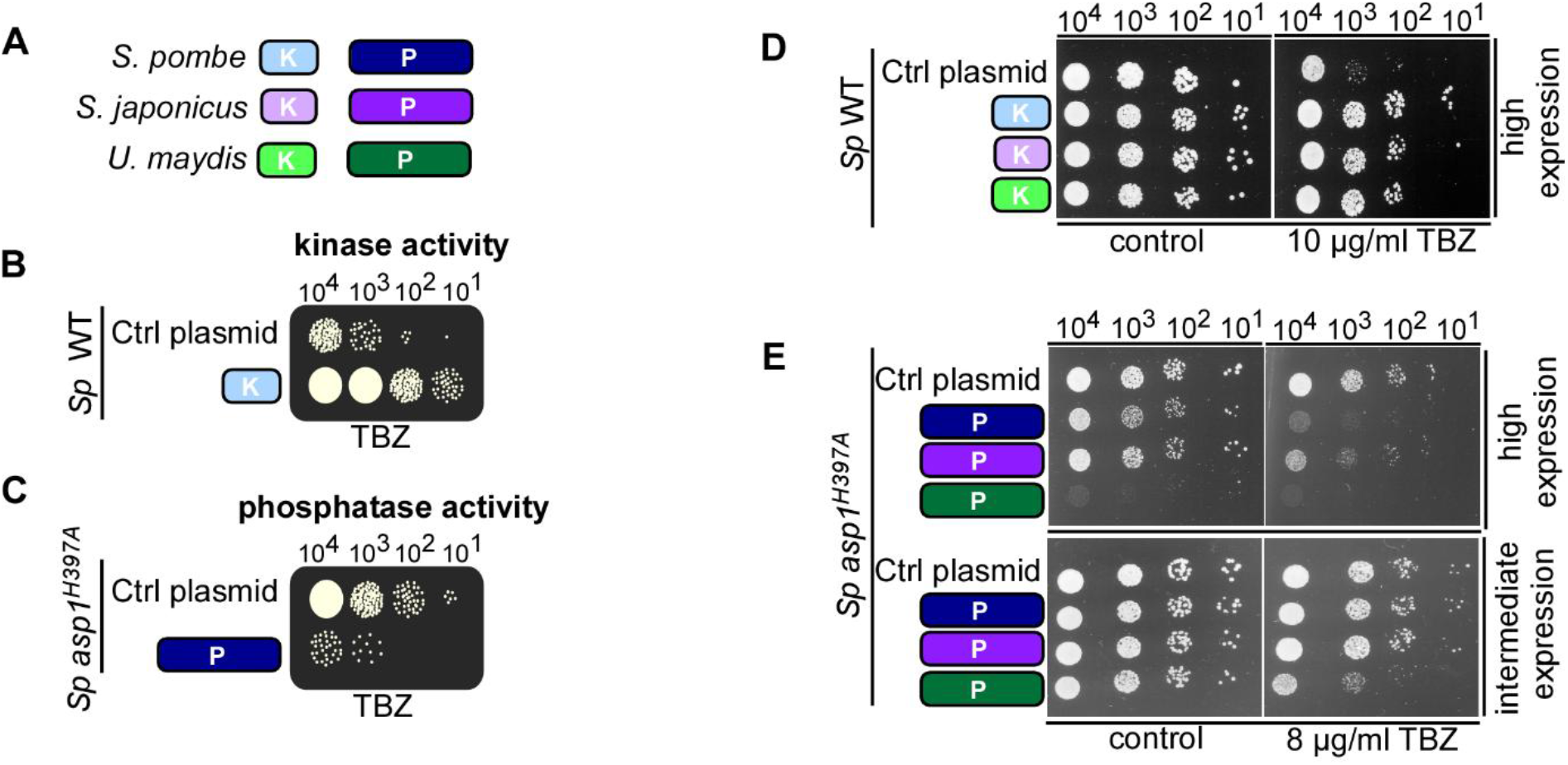
Determination of Asp1 kinase and phosphatase activities using an *S. pombe in vivo* read-out system. **(A)** Schematic representation of the Asp1 variants used: *S. pombe* kinase (light blue, amino acids 1-364) and phosphatase (dark blue, amino acids 365-920), *S. japonicus* kinase (light purple, amino acids 3-366) and phosphatase (dark purple, amino acids 367-925) and *U. maydis* kinase (light green, amino acids 1-336) and phosphatase (dark green, amino acids 337-922). **(B)** Diagrammatic representation of a serial dilution patch test (10^4^, 10^3^, 10^2^ and 10 cells) as an *in vivo* read-out of PPIP5K kinase function in *Sp* WT cells under *nmt1^+^* promoter derepressing (high expression) conditions on plates containing Thiabendazole (TBZ). (**C**) Diagrammatic representation of a serial dilution patch test (10^4^, 10^3^, 10^2^ and 10 cells) as an *in vivo* read-out of PPIP5K phosphatase function in the *Sp asp1^H397A^* strain under *nmt1^+^*promoter derepressing (high expression) conditions on plates containing TBZ. (**D**) Serial dilution patch test (10^4^, 10^3^, 10^2^ and 10^1^ cells) of a *Sp* WT strain expressing plasmid-encoded kinase (K) domains from *S. pombe*, *S. japonicus* or *U. maydis* or carrying a control plasmid (ctrl) on plates without (control) or with 10 µg/ml TBZ. Cells were grown on media without thiamine which fully derepresses the *nmt1^+^*promoter. Plates were incubated at 25 °C for 7 days. (**E**) Serial dilution patch test (10^4^, 10^3^, 10^2^ and 10 cells) of an *Sp asp1^H397A^* strain expressing plasmid-encoded phosphatase (P) domains from *S. pombe*, *S. japonicus* or *U. maydis* or carrying a control plasmid (ctrl) on plates without (control) or with 8 µg/ml TBZ. High expression: top panels, medium without thiamine. Intermediate expression: bottom panels, medium with 0.05 µM thiamine [37]. Plates were incubated at 25 °C for 5 days.

### SjAsp1 restricts the yeast-to-hypha transition in *S. japonicus*

Having established conserved enzymatic activity, we asked whether the biological output of PPIP5K signalling was similarly conserved. PPIP5K activity promotes the yeast-to-pseudohyphal transition in *S. pombe* and filamentous growth in *U. maydis* [23, 24]. We therefore examined invasive and hyphal growth in *S. japonicus*. Substrate invasion of *S. japonicus* cells necessitates the switch from yeast surface to invasive hyphal growth [38]. Thus, we analysed the ability of *Sj* WT and *Sj asp1Δ* strains to invade a substrate by growing these strains on solid media (Fig. 4A, top panels) followed by washing and rubbing off cells/colonies present on the surface of the agar (Fig. 4A, bottom panels). Invasive colonies were counted microscopically. Surprisingly, we found that the *Sj asp1Δ* strain formed significantly more invasive colonies than the wild-type strain (Fig. 4A, right panel). Next, we analysed mycelial expansion of the two *S. japonicus* strains by patching cells from log-phase growth onto YEMA plates (Fig. 4B)[38]. The area of mycelial expansion was greater in the mutant than in the wild-type strain (Fig. 4B, right diagram).

**Fig. 4.**
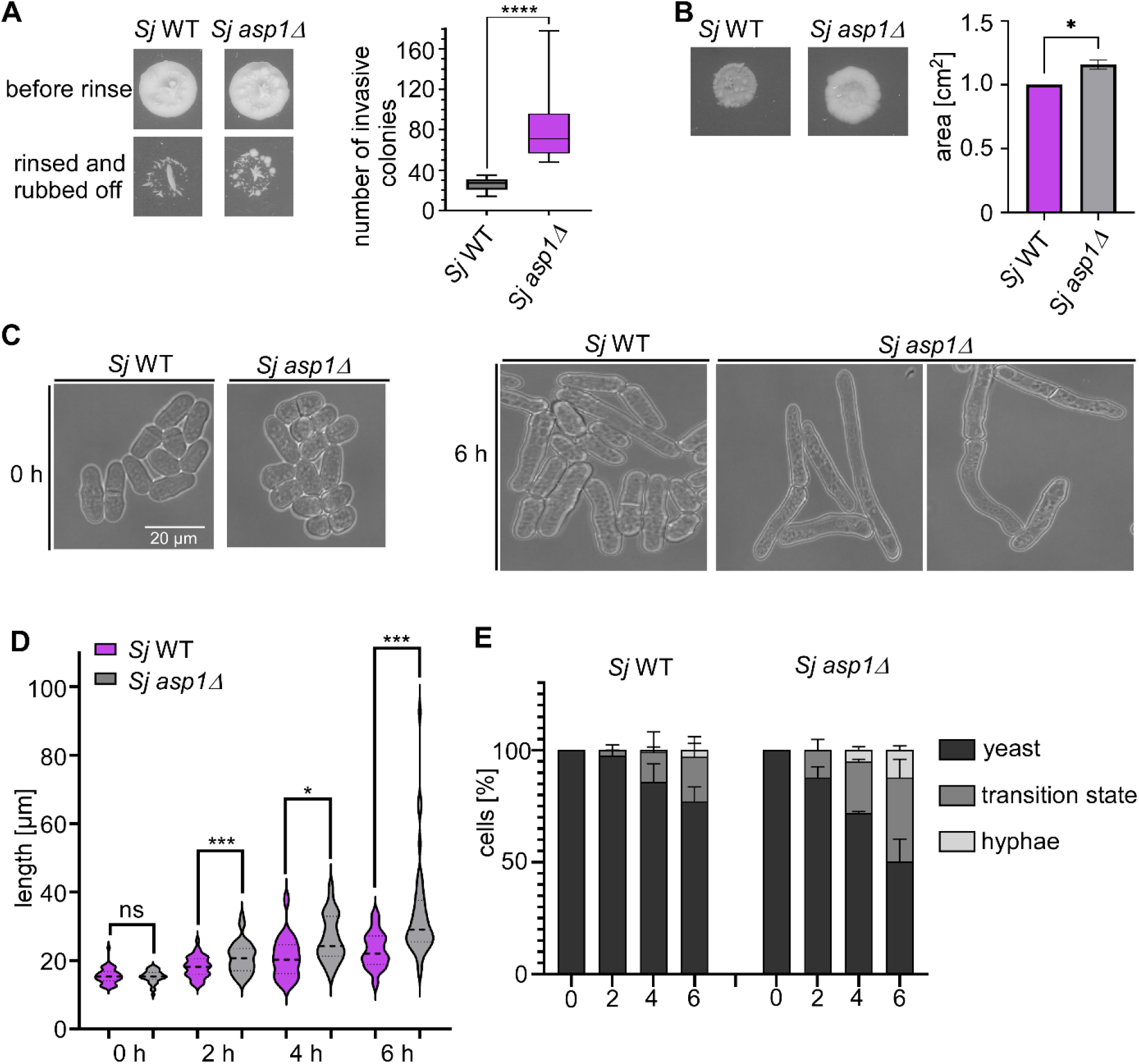
Absence of *S. japonicus* Asp1 accelerates the switch to hyphal growth. **(A)** Macroscopic image of *S. japonicus* growth of WT and *asp1Δ* strains on YE5S plates after 4 days at 30 °C before rinsing (top panel) and after rinsing (bottom panel). Right: Quantification of invasively grown colonies. The data represent three experiments, each with three biological replicates. Statistical comparison was performed using a Kolmogorov–Smirnov test. ****, *P*<0.0001. **(B)** Mycelial expansion of *Sj* WT and *Sj asp1Δ* cells: 10^5^ cells were plated onto YEMA plates and incubated for 4 days at 30 °C. Left: Representative images of cell migration. Right: Quantification of colony area. The data represent three datasets with 18 colonies. The statistical analysis was performed using Welch’s test. *, *P* < 0.05; **(C)** Brightfield images of WT and *Sj asp1Δ* cells at timepoint 0- and 6-hours incubation with CPT (0.2 µM). Scale bar: 20 µm **(D)** Measurement of cell length before and after CPT treatment at the times indicated. Data represent three independent experiments. Data from all experiments were pooled into a single graph (n = 70–101 cells per strain and condition). Statistical comparisons were performed using two-tailed Welch’s t-tests. ns, not significant; *, *P*< 0.05; ***, *P*< 0.001 **(E)** Quantification of yeast-to-hypha transition. Data represent three independent experiments (n = 10–41 per repetition).

To determine if SjAsp1 affected early transition events required for hyphal development, the two *S. japonicus* strains were incubated with low doses of Camptothecin (CPT), a DNA topoisomerase inhibitor, which quickly induces hyphal differentiation [39]. The yeast to hyphal transition occurs in the following way: yeast cells will progress to cells with many vacuoles that will eventually be localized at one cell end (transition state), followed by vacuole fusion and hyphal formation (hyphal state) [40]. Cells become highly elongated during morphogenesis. We analysed growth alterations in CPT containing liquid medium for 6 hrs and found that cell elongation, appearance of elongated cells with multiple vacuoles (here named transition stage) and appearance of hyphal cells occurred earlier and with a higher frequency in the *Sj asp1Δ* strain than in the WT strain (Fig. 4C-E). Thus, Asp1 in *S. japonicus* negatively affects the yeast to hyphal transition.

### SjAsp1 promotes adhesion and invasive pseudohyphal growth when expressed in *S. pombe*

The opposing functions of SpAsp1 and SjAsp1 raised a central question: is the direction of Asp1-dependent regulation encoded by the Asp1 protein itself, or by the cellular environment in which its products act? For *S. pombe*, invasive growth requires the adhesion of yeast cells to the agar surface followed by invasive pseudohyphal growth into the agar. To answer this question, we expressed full-length, kinase-domain or phosphatase-domain variants of SpAsp1 and SjAsp1 in *S. pombe* wild-type (WT) and *asp1^H397A^* cells and quantified invasive growth (Fig. 5A).

**Fig. 5.**
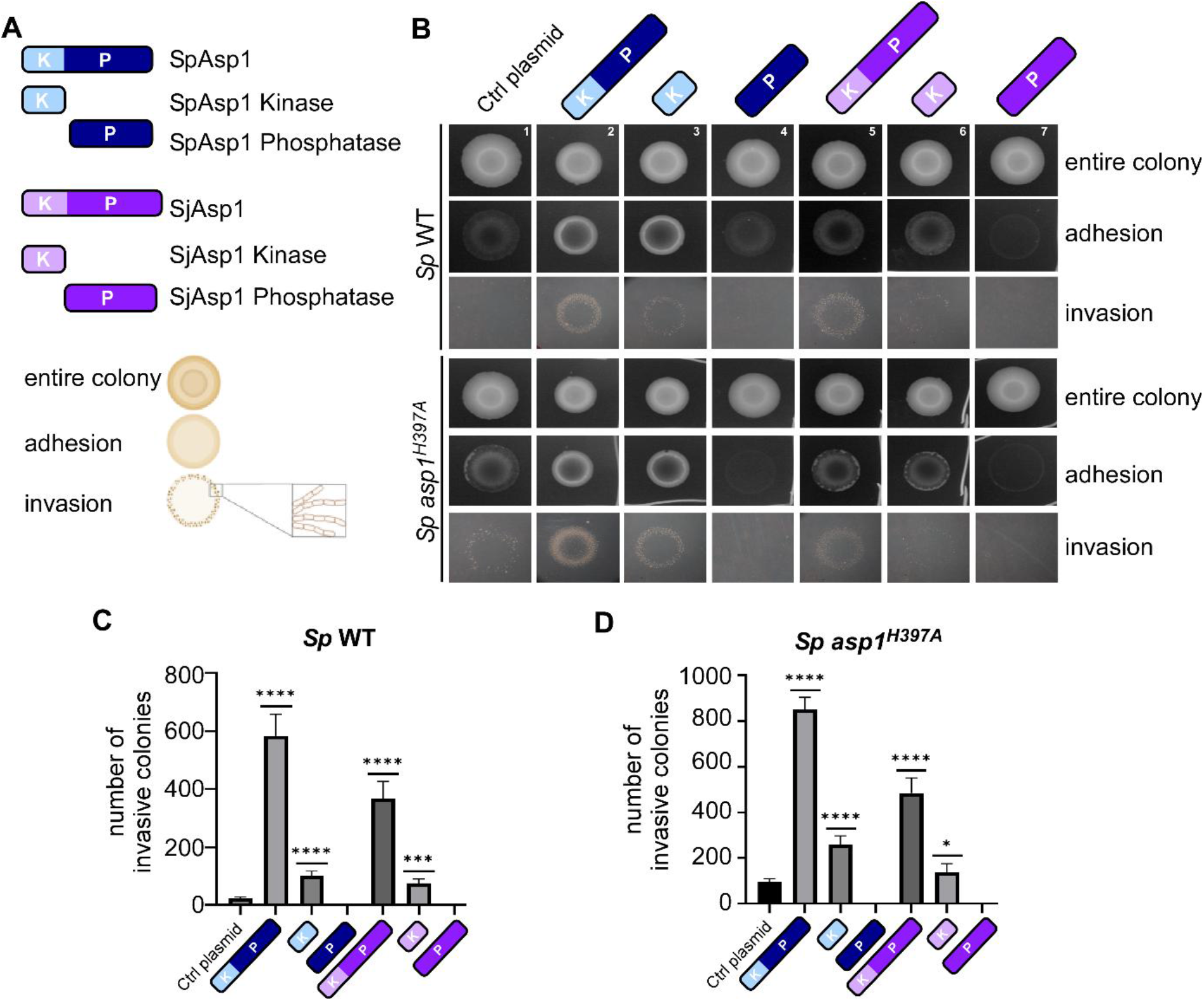
SjAsp1 phenocopies SpAsp1 when expressed in *S. pombe*. (**A**) Schematic representation of the plasmid-expressed Asp1 variants used in this experiment. SpAsp1 full-length (light and dark blue, amino acids 1-920), SpAsp1 kinase (light blue, amino acids 1-364), SpAsp1 phosphatase (dark blue, amino acids 365-920), SjAsp1 full-length (light and dark purple, amino acids 3-925), SjAsp1 kinase (light purple, amino acids 3-366) and SjAsp1 phosphatase (dark purple, 367-925), top panel. Bottom panels: Schematic overview of the invasive growth assay setup in *S. pombe*. (**B**) Representative images of adhesive and invasive colony morphology in *Sp* WT and *Sp asp1^H397A^* strains expressing plasmid-borne full-length, kinase or phosphatase Asp1 variants from *S. pombe* and *S. japonicus*. (**C+D**) **C**: Quantification of the number of invasive colonies in the *Sp* WT transformants. Data represent six biological replicates. **D**: Quantification of the number of invasive colonies in the *Sp asp1^H397A^* strain. Data represent six biological replicates. Statistical comparisons were performed using multiple two-tailed Welch’s t-tests. *, *P* < 0.05; ***, *P* <0.001; ****, *P* < 0.0001.

Intracellular 1,5(PP)_2_-InsP_4_ levels dose-dependently regulate both adhesion and invasion in *S. pombe*: higher than wild-type 1,5(PP)_2_-InsP_4_ levels will lead to increase adhesion and invasion while lower than wild-type levels have the opposite effect [24]. Indeed, increasing intracellular 1,5(PP)_2_-InsP_4_ by expressing full-length SpAsp1 or the SpAsp1 kinase in WT and *asp1^H397A^* transformants led to increased adhesion and invasive growth. (Fig. 5B panels 2,3; quantification in Fig. 5C and D). As the *asp1^H397A^* strain has more 1,5(PP)_2_-InsP_4_ than the WT strain, invasive growth of this strain is *per se* increased compared to the WT [21, 24]. Interestingly, expression of SjAsp1 full-length or kinase domain had a similar effect namely an increase in the number of invasively growing colonies (Fig. 5B, panels 5 and 6; quantification in Fig 5C and 5D). Furthermore, expression of the SpAsp1 phosphatase or SjAsp1 phosphatase domains resulted in an inability of the transformants to adhere to the substrate and grow invasively (Fig. 5 B, panels 4 and 7; quantification in 5C-D). Thus, although SjAsp1 restricts the yeast-to-hyphal transition in *S. japonicus*, it promotes adhesion and invasive pseudohyphal growth when expressed in *S. pombe*. The direction of the PPIP5K-dependent response is therefore imposed by the species-specific cellular context rather than by an intrinsic activating or inhibitory property of the SjAsp1 protein.

### Conserved PPIP5K activity produces species-specific microtubule and mitochondrial outputs

Because yeast-to-hyphal growth depends on coordinated cytoskeletal organization, we next asked whether the context-dependent effect of Asp1 on this transition was accompanied by divergence in cytoskeletal regulation.

In *S. pombe* an intact actin cytoskeleton is a prerequisite for the yeast to pseudohyphal transition while altered interphase microtubule (MT) dynamics can promote this process [24]. In *U. maydis* and *S. japonicus* the actin cytoskeleton plays a major role in the yeast to hypha transition [40, 41]. Thus, we tested if absence of the respective Asp1 protein affected the MT and actin cytoskeletons of the three fungi.

Asp1 in *S. pombe* and *U. maydis* modulates MT organization and dynamics [23] and loss of the respective Asp1 encoding ORF resulted in cells with increased sensitivity towards the MT poison TBZ (Fig. 6A) [23]. This was also the case for *S. japonicus* for the *asp1Δ* strain (Fig. 6B). To determine if *Sj asp1Δ* cells also showed aberrant interphase MT organization as we had previously determined for *U. maydis* and *S. pombe asp1Δ* strains [23], we analyzed the spatial distribution of interphase MTs in *Sj* WT and *asp1Δ* cells. In both *S. japonicus* var. *versatilis* and *S. japonicus*, the interphase MT cytoskeleton has been described as consisting of longitudinal bundles oriented parallel to the long axis of the cell [40, 42, 43] and this type of spatial distribution was found for our *Sj* WT and *asp1Δ* strains (Fig. 6C). We therefore analyzed MT organization in interphase cells of *Sj* WT and *asp1Δ* strains by investigating the orientation of MTs along the cell’s longitudinal axis (MT coherency). Using this approach, we found no significant difference in MT coherency between WT and *asp1Δ* cells (Fig. 6D). We next compared an MT-linked organelle output. In *S. pombe*, mitochondria associate with interphase MTs, and loss of SpAsp1 kinase activity disrupts this interaction and causes a strongly aberrant mitochondrial morphology (Fig 6E; quantification in 6H, further explanation of the wild-type and aberrant phenotypes in Fig. S2) [21, 44–46].*Um asp1Δ* cells likewise showed a significant increase in aberrant mitochondrial morphology (Fig. 6G; quantification in 6H). By contrast, mitochondrial organization was not statistically altered in *Sj asp1Δ* cells (Fig. 6F; quantification in 6H), despite the reported association between MTs and mitochondria in *S. japonicus var. versatilis* [47]. These findings retain MT regulation as a conserved branch of fungal PPIP5K biology, but show that its cellular consequences are species-specific: Asp1 loss affects mitochondrial organization in *S. pombe* and *U. maydis*, whereas this output appears to be uncoupled from Asp1 in *S. japonicus*.

**Fig. 6.**
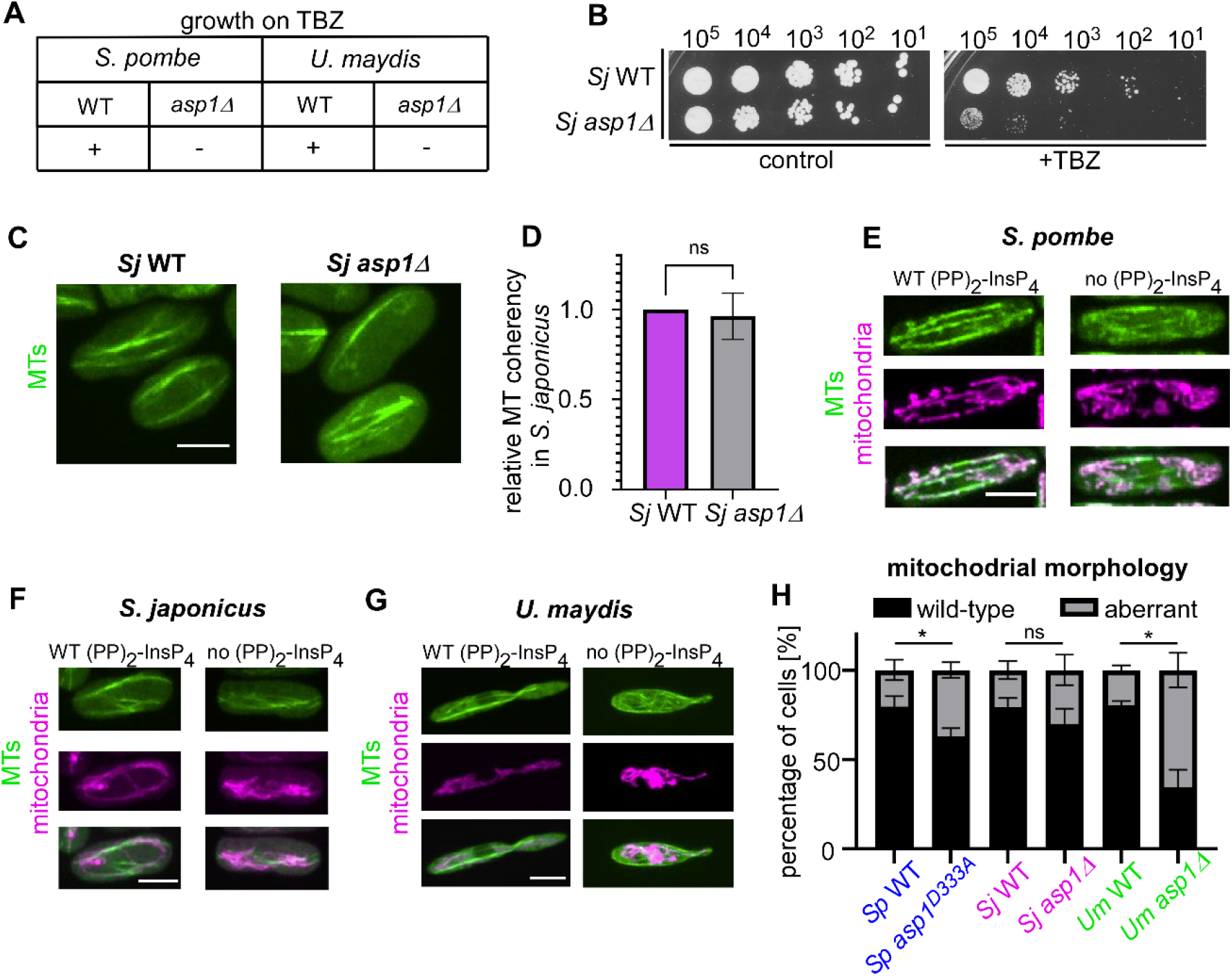
1,5(PP)_2_-InsP_4_ is required for mitochondrial network architecture in *S. pombe* and *U. maydis*, but not in *S. japonicus*. (**A**) Schematic summary of the growth phenotypes of *S. pombe* and *U. maydis* WT and *asp1Δ* cell adapted from [23, 24]. (**B**) Serial dilution patch test (10^5^-10^1^ cells) of *S. japonicus* WT and *asp1Δ* cells grown on YE5S plates (control, left) or YE5S plates supplemented with 8 µg/ml TBZ (right). (**C**) Fluorescence microscopy of fixed *Sj* WT and *asp1Δ* cells expressing Atb2-GFP. Fluorescence intensity was not statistically altered (mean gray value normalized to WT: 1:1.29, p=0.26). (**D**) Quantification of MT organization *in S. japonicus*. MT orientation was analyzed using the ImageJ OrientationJ plugin, which measures the degree to which MTs are aligned with the longitudinal axis of the cell. Higher coherency values indicate a more ordered MT array, whereas lower values reflect increased MT disorganization. Relative MT coherency in *Sj asp1Δ* cells normalized to the Sj WT control (right). Data are derived from three independent experiments comprising a total of 60 WT and 62 *asp1Δ* cells. Statistical significance was assessed using Welch’s *t*-test; ns, not significant. (**E**) Live-cell fluorescence microscopy of *Sp* WT and *asp1^D333A^* (no 1,5(PP)_2-_InsP_4_) cells expressing Atb2-GFP (α-tubulin, green) and Cox4-RFP (complex IV of the inner mitochondrial membrane, red). Bottom image shows merged image. (**F**) Fluorescence microscopy of fixed *Sj* WT and *asp1Δ* (no 1,5(PP)_2-_InsP_4_) cells expressing Atb2-GFP (α-tubulin, green). Mitochondria were stained with MitoTracker CMXRos (red) prior to fixation. Bottom image shows merged image. (**G**) Live-cell fluorescence microscopy of *Um* WT and *asp1Δ* (no 1,5(PP)_2-_InsP_4_) cells expressing Tub1-GFP (α-tubulin, green). Mitochondria were stained with TMRE (red). Bottom image shows merged image. (**H**) Quantification of mitochondrial phenotypes of *Sp*, *Sj* and *Um* WT and *asp1Δ* strains. Data are derived from three independent experiments. *S. pombe*: WT: n=187, *asp1^D333A^*: n=173, *S. japonicus*: WT: n=163, *asp1Δ*: 85 cells, *U. maydis*: WT: n= 153, *asp1Δ*: n=105. Statistical analysis was performed using a Welch’s test. ns: not significant *, *P* < 0.05. Scale bars: 5 µm.

### Loss of SjAsp1 confers resistance to Arp2/3 inhibition and alters cortical actin patch organization

After having analysed the if MT-organization in *S. japonicus asp1Δ cells was* significantly altered, we next analysed the actin cytoskeleton, which is essential for the *S. japonicus* yeast-to-hypha transition [40]. Asp1 has previously been linked to Arp2/3 function in *S. pombe*, where *Sp asp1Δ* cells exhibit Arp2/3-related defects [19]. The presence of CK666, a small-molecule inhibitor of the Arp2/3 complex [48], led to reduced growth of *Sp asp1Δ* and *Um asp1Δ* cells compared to the corresponding wild-type cells (Fig. S3), thus also supporting a role of UmAsp1 in Arp2/3 modulation. However, and in contrast, the opposite phenotype was observed for *Sj asp1Δ* cells as these were highly resistant to CK666 (Fig. 7A). Thus, we analysed cortical actin patch organization in the two *S. japonicus* strains using LifeAct-GFP [40]. A LifeAct-GFP-expressing *Sj asp1Δ* strain was generated and imaged alongside a *Sj* WT strain expressing LifeAct-GFP and α-tubulin Atb2-mCherry. Live-cell imaging of both strains was done on the same microscope slide. As the WT expressed Atb2-mCherry, we could easily identify the cells of the two strains (Fig. 7B, left panel). The number of actin patches increases approximately proportionally with cell length [49], and thus, the number of actin patches per cell area were determined. Mean patch density was 0.49 ± 0.09 patches/µm² for *Sj* WT cells and 0.72 ± 0.14 patches/µm² for *Sj asp1Δ* cells. This corresponds to a highly significant increase of approximately 47% in the mutant (Fig. 7C). We next examined the properties of individual actin patches such as the size of the patches and their fluorescence signal intensity. Mean patch area increased significantly from 0.21 ± 0.08 µm² in *Sj* WT to 0.24 ± 0.11 µm² in *Sj asp1Δ* cells (Fig. 7D). Furthermore, LifeAct-GFP-labelled actin patches in *Sj asp1Δ* cells were brighter than those in *Sj* WT cells: mean fluorescence intensity increased from 4.97 ± 3.31 arbitrary units (a.u.) in *Sj* WT cells to 8.37 ± 6.01 a.u. in *Sj asp1Δ* cells, corresponding to a 68% increase in F-actin fluorescence (Fig. 7E). To determine whether loss of SjAsp1 affects actin patch lifetime, individual patches were followed by time-lapse fluorescence microscopy. Mean patch lifetime was similar: 15.28 ± 2.93 s in *Sj* WT and 15.35 ± 3.33 s in *Sj asp1Δ* cells (Fig. 7F), indicating that the increased patch density/area in *Sj asp1Δ* cells was not caused by prolonged patch lifetime. During live-cell imaging, we observed that cortical actin patches in *Sj asp1Δ* cells were less restricted to the cell ends, with an increased abundance of patches in the cell middle compared with *Sj* WT cells. Our quantification showed that the mean number of patches in the cell middle increased from 8.78 ± 4.22 in *Sj* WT to 16.71 ± 6.55 in *Sj asp1Δ* cells, corresponding to an approximately 90% increase in the mutant (Fig. 7G). Thus, loss of SjAsp1 results in increased actin patch density/area, increased patch-associated F-actin fluorescence, and an increased abundance of patches in the cell middle, while patch lifetime remains unchanged. CK666 inhibits Arp2/3-dependent actin assembly and thereby interferes with the formation of cortical actin patches. The pronounced resistance of *Sj asp1Δ* cells to CK666, together with the observed increase in patch density and F-actin fluorescence, shows that SjAsp1 negatively impacts on Arp2/3-dependent cortical actin patches.

**Fig. 7.**
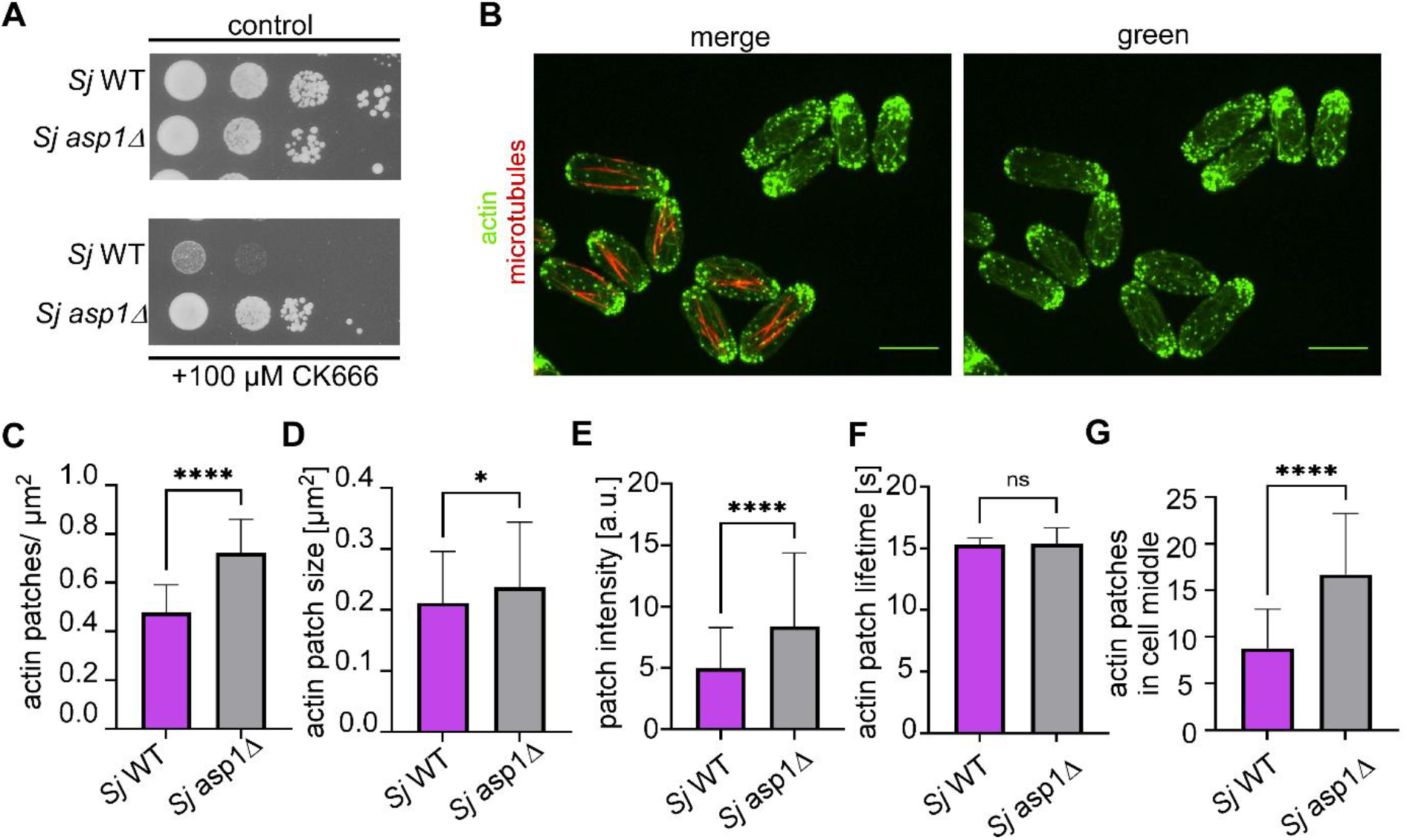
SjAsp1 restricts Arp2/3-dependent actin assembly. **(A)** Serial dilution patch test assay (10^4^ to 10^1^ cells) of *Sj* WT and *Sj asp1*Δ strains on YE5S plates supplemented with DMSO (control) or the Arp2/3 inhibitor CK666 (100 µM). Plates were incubated for 3 days at 25 °C. **(B)** Representative fluorescence microscopy images of *Sj* WT and *Sj asp1Δ* cells expressing LifeAct-GFP to visualize F-actin (green). *Sj* WT cells additionally express Atb2-mCherry (α-tubulin, red), allowing WT cells to be distinguished from *asp1Δ* cells when imaged on the same slide. The left panel shows the merged LifeAct-GFP and Atb2-mCherry channels, whereas the right panel shows the LifeAct-GFP channel only. Scale bars, 10 µm. **(C)** Quantification of actin patch density/cell area (patches/µm²). Patch density was calculated by dividing the number of actin patches by the cell area. 45 cells in *Sj* WT and 42 cells in *Sj asp1Δ* cells (3 biological replicates/strain) were analysed. Mean actin patch density was 0.49 ± 0.09 patches/µm² for *Sj* WT and 0.72 ± 0.14 patches/µm² for *Sj asp1Δ* (mean ± SD; Mann–Whitney U test, ****, *p* < 0.0001). **(D)** Quantification of actin patch size (µm²) using Fiji/ImageJ. 195 actin patches for *Sj* WT and 204 actin patches for *Sj asp1Δ* were analysed (3 biological replicates/strain). Mean actin patch size was 0.21 ± 0.08 µm² for *Sj* WT and 0.24 ± 0.11 µm² for *Sj asp1Δ* (mean ± SD; Mann–Whitney U test, *, *p* = 0.0213). **(E)** Quantification of actin patch fluorescence intensity using Fiji/ImageJ. 195 actin patches for *Sj* WT and 204 actin patches for *Sj asp1Δ* (3 independent biological replicates/strain) were analysed. Mean actin patch intensity was 4.97 ± 3.31 a.u. for *Sj* WT and 8.37 ± 6.01 a.u. for *Sj asp1Δ* (mean ± SD; Mann–Whitney U test, ****, *p* < 0.0001). **(F)** Quantification of actin patch lifetime. Patch lifetime was determined via time-lapse fluorescence microscopy by measuring the time between the appearance and disappearance of an individual actin patch. 60 actin patches were analysed for each strain (3 independent biological replicates/strain). Mean actin patch lifetime was 15.28 ± 2.93 s for *Sj* WT and 15.35 ± 3.33 s for *Sj asp1Δ* (mean ± SD; unpaired Welch’s *t*-test, ns=not significant, *p* = 0.9401). **(G)** Quantification of the spatial distribution of actin patches in the middle of the cell, which was divided into 3 parts. 45 cells in *Sj* WT and 42 cells in *Sj asp1Δ* cells (3 biological replicates/strain) were analysed. Patches in cell middle: 8.78 ± 4.22 in *Sj* WT and 16.71± 6.55 in *Sj asp1Δ* (mean ± SD; Mann–Whitney U test, ****, *p* < 0.0001).

### The *S. pombe* Arp2/3 complex is a target of inositol 1,5(PP)_2_-InsP_4_

Having established that species context dictates whether Asp1 functions as a positive or negative modulator of actin-driven morphogenesis, we sought the direct biochemical mechanism underlying this signaling output. In *S. pombe*, *Asp1* was originally identified through genetic suppression of Arp2/3 mutants, yet the Asp1 protein itself did not co-fractionate with the Arp2/3 complex [19]. This suggests that the signaling output might not be mediated by physical interactions with the enzyme itself, but rather through its catalytic products. 5PP-InsP_5_ and 1,5(PP)_2_-InsP_4_, have been proposed to regulate protein function directly by interacting with protein targets [7] and via protein pyrophosphorylation [15]. Thus, to identify potential InsPP targets in *S. pombe* possibly including actin modulators, we incubated *S. pombe* cell lysates with affinity reagent biotin-1,5(PCP)_2_-InsP_4_ (b-1,5(PCP)_2_-InsP_4_) (Fig. 8A). In this reagent, the pyrophosphate moieties have been replaced with non-hydrolyzable bisphosphonate groups (PCP-groups) and the PP-InsP analog is derivatized at the 3-position with biotin via a soluble linker, to enable immobilization onto streptavidin-coated sepharose beads. In addition, immobilized b-5PCP-InsP_5_, immobilized b-InsP_6_, and unmodified streptavidin-coated beads were used [50–52]. After incubation and washing, interacting proteins were competitively eluted by adding an excess of the corresponding non-biotinylated PP-Ins/InsP species (diagrammatic representation of experimental set-up shown in Fig. 8B). Eluates were analysed via MS-based quantitative proteomics.

**Fig. 8.**
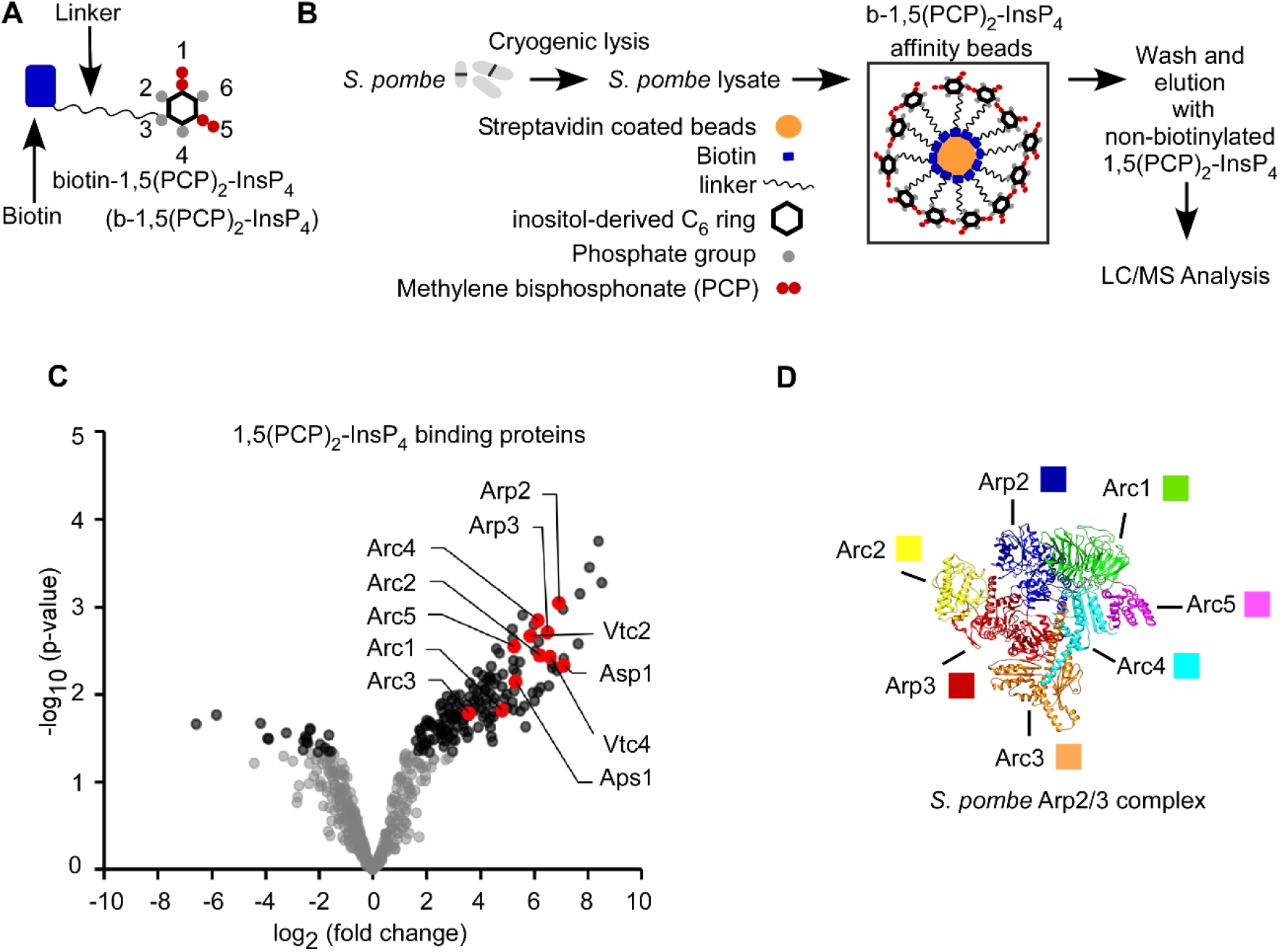
Identification of *S. pombe* Arp2/3 complex as an 1,5(PP)_2_-InsP_4_-binding protein complex. **(A)** Schematic representation of the biotinylated, non-hydrolysable 1,5(PCP)_2_-InsP_4_ affinity reagent b-1,5(PCP)_2_-InsP_4_ [52]. **(B)** Experimental workflow for the identification of 1,5(PP)_2_-InsP_4_-binding proteins. Cryogenically lysed *S. pombe* cells were incubated with b-1,5(PCP)_2_-InsP_4_-loaded streptavidin-coated Sepharose beads. After washing, bound proteins were competitively eluted using excess non-biotinylated 1,5(PCP)_2_-InsP_4_ and identified by quantitative LC-MS-based proteomics. **(C)** Volcano plot showing proteins enriched on b-1,5(PCP)_2_-InsP_4_-coated beads relative to control beads. The x-axis shows the log₂ fold change in protein abundance averaged over the replicates, and the y-axis the −log₁₀ of the student’s t-test *p*-value as means for visualization of statistical significance. The statistical significance cut-off was set at 5% permutation-based false discovery rate (5% FDR: grey/black datapoint regions) using the significance analysis of microarrays (SAM) method. Proteins known to be regulated by 1,5(PP)_2_-InsP_4_ (Vtc2 and Vtc4) and enzymes involved in 1,5(PP)_2_-InsP_4_ levels (Asp1 and Aps1) [7, 21, 55, 56], together with all seven subunits of the Arp2/3 complex (Arp2, Arp3 and Arc1–Arc5), are highlighted as red datapoints and labelled. **(D)** Structural representation of the *S. pombe* Arp2/3 complex (PDB ID: 6W17). Individual subunits are colour-coded as follows: Arp2 (blue), Arp3 (red), Arc1 (green), Arc2 (yellow), Arc3 (orange), Arc4 (turquoise), and Arc5 (magenta). The bound nucleation-promoting factor and actin molecules were omitted for clarity. The structure was visualized using UCSF Chimera version 1.18 [57].

Mass spectrometry (MS)-based analysis of pulldowns of b-1,5(PCP)_2_-InsP_4_-coated beads vs. control beads identified several proteins expected to associate with 1,5(PP)_2_-InsP_4_ (Fig.8C). These included the 1,5(PP)_2_-InsP_4_-regulated, SPX-harbouring proteins Vtc2 and Vtc4 [13, 25, 27, 53, 54], and the key enzymes involved in maintaining 1,5(PP)_2_-InsP_4_ levels, the inositol pyrophosphatase Aps1 [55] and Asp1 [21, 56]. Notably, these proteins were among the most highly enriched proteins identified in the MS analysis. Their recovery provides validation of the affinity-enrichment approach and supports the physiological relevance of the identified 1,5(PP)_2_-InsP_4_-associated protein candidates.

Remarkably, all 7 components of the highly conserved Arp2/3 complex Arp2, Arp3, Arc1, Arc2, Arc3, Arc4, and Arc5 were among the most enriched proteins identified (Fig. 8C-D). Although this analysis cannot distinguish if one/several Arp2/3 members are associated with1,5(PP)_2_-InsP_4_, it establishes a a long-sought biochemical bridge between inositol pyrophosphate signaling and the core actin nucleation machinery. Analysis of 5PCP-InsP_5_ and InsP_6_ protein targets uncovered a similar pattern of protein interactions to that observed with 1,5(PCP)_2_-InsP_4_, including all seven Arp2/3 complex subunits together with Vtc2, Vtc4, Asp1 and Aps1 (Fig.S4), as has been shown for previous pull-down experiments utilizing these reagents [52].

## Discussion

### *S. japonicus* shows a distinct PPIP5K output in our three-species comparison

Our comparative analysis separates conservation of PPIP5K enzymatic activity from conservation of its biological output. The Asp1 homologs of the ascomycetes *Schizosaccharomyces pombe* and *Schizosaccharomyces japonicus* and the basidiomycete *Ustilago maydis* retain a conserved kinase-phosphatase architecture and are required for detectable production of 1,5-(PP)₂-InsP₄ in their native organisms [21]. Absence of Asp1 in all three organisms results in elevated PP-InsP_5_ levels, but in the *Um asp1Δ* strain, this was extremely high. Such an increase has not been observed for any other studied eukaryote and raises the intriguing possibility that in this basidiomycete, the PPIP5K enzyme/its product(s) act as negative regulators of upstream enzymes responsible for PP-InsP_5_ synthesis.

Although enzyme architecture and function are similar for the three PPIP5K family members, their modulation of specific biological processes differ. Asp1 is required for mitochondrial organization in *S. pombe* and *U. maydis* but not in *S. japonicus.* Strikingly, Asp1 has opposing effects on morphogenesis and the response to Arp2/3 inhibition: in *S. pombe* and *U. maydis*, Asp1 promotes invasive pseudohyphal and wild-type hyphal growth, respectively, and its loss increases sensitivity to the Arp2/3 inhibitor CK666. However, in *S. japonicus*, Asp1 restricts the yeast-to-hypha transition, whereas *asp1* deletion confers CK666 resistance. Thus, the closely related fission yeasts display opposing outputs, while the distantly related *S. pombe* and *U. maydis* behave similarly.

Cross-species expression demonstrates that this inversion is not an intrinsic property of the SjAsp1 protein. In *S. pombe*, SjAsp1 and its catalytic domains reproduce the morphogenetic effects of SpAsp1, and SjAsp1 also increases CK666 resistance when expressed in *S. pombe*. SjAsp1 therefore adopts both the morphogenetic and Arp2/3-related outputs of the recipient species. Together with the established dose-dependent control of *S. pombe* invasive growth by Asp1 catalytic activity [21, 24] these findings indicate that SjAsp1 supplies a functional inositol pyrophosphate signal, whereas the host network determines its biological consequence.

Only a few other studies connect PPIP5K-family proteins with the promotion or correct organization of fungal morphogenesis. The *Aspergillus nidulans* homologue VipA contributes to growth-zone selection and normal hyphal organization [23]. Furthermore, two studies of the morphogenetic function of the *Saccharomyces cerevisiae* PPIP5K member Vip1 exist, but they reached conflicting conclusions regarding its effect on invasive and pseudohyphal growth [24, 58]. Apart from this unresolved case, SjAsp1 therefore represents the clearest fungal example in which a PPIP5K restricts a morphogenetic transition.

The shared behavior of *S. pombe* and *U. maydis* excludes several simple explanations for the inversion. It cannot be attributed to phylogenetic distance, because the shared output spans the ascomycete and basidiomycete lineages. Nor can it be explained by the existence or extent of a dimorphic transition, because *U. maydis* undergoes a pronounced switch to polarized hyphal growth yet retains the *S. pombe*-type Asp1 output. The results instead point to a specific reassignment of PPIP5K signalling within the *S. japonicus* cellular network.

### A conserved signal can acquire an opposite biological output

The *S. japonicus* phenotype represents a reversal rather than a quantitative weakening of Asp1 function. SjAsp1 restricts morphogenesis in its native species but promotes invasive growth when expressed in *S. pombe*. Because the same enzyme adopts the morphogenetic output of the recipient species, the inversion is most likely located downstream of signal production.

The output of a conserved signaling module can be reversed by changes in its regulatory context. For example, it has been shown that rewiring feedback around the same MAPK-ERK module altered and reversed the resulting cellular response [59]. Thus, signal identity alone does not determine biological outcome.

Comparable divergence of fungal morphogenesis has been described for the budding yeasts *Saccharomyces cerevisiae* and members of the *Saccharomyces bayanus* species complex; lineages separated by approximately 20 million years [60]. Here, increasing or decreasing cAMP-PKA activity had opposite effects on filamentous growth and this divergence extended across several downstream pathway components, indicating substantial reorganization of the morphogenesis-signalling network [61]. Species-specific adaptation of a conserved regulatory system was also recently described for membrane homeostasis in *S. pombe* and *S. japonicus*, where the Mga2-Ole1 pathway is tuned differently to the distinct lipid environment of the two species [62]. Furthermore, opposite effects of the same stress-response pathway have been observed for *S. pombe* and *S. japonicus*. Sty1 is a stress-activated mitogen-activated protein kinase that coordinates cellular adaptation to adverse conditions. When the actin cytoskeleton is perturbed in *S. pombe*, Sty1 limits formation of the actomyosin ring that drives cytokinesis. In *S. japonicus*, however, Sty1 activity is needed for actomyosin ring formation [63]. The reason for this difference is unresolved, but the authors suggested that it may be related to the distinct timing of ring formation in the two species [63].

### Arp2/3 is a candidate effector of 1,5-(PP)_2_-InsP_4_

Genetic evidence first linked the PPIP5K/Asp1/Vip1 family to Arp2/3-dependent actin organization. In *S. pombe*, increased *asp1^+^* dosage suppressed an *arp3* mutant, whereas *asp1* deletion was synthetically lethal with mutations in Arp2/3 subunits and disrupted cortical actin-patch localization [19]. In *S. cerevisiae*, deletion of the Asp1 homologue VIP1 caused a synthetic growth defect with loss of LAS17, encoding the WASP-family activator of Arp2/3, and rescue required Vip1 kinase activity, linking PPIP5K-generated inositol pyrophosphates to the Arp2/3 pathway [56].

Our recovery of all seven *S. pombe* Arp2/3 subunits with biotin-1,5-(PCP)₂-InsP₄ provides a possible biochemical basis for these genetic observations. Their concerted enrichment suggests capture of an assembled complex through association of one or more components with the reagent. However, the experiment does not establish direct binding, identify the interacting subunit, or show that ligand association alters Arp2/3 activity. Nevertheless, this data gives a possible explanation for the finding more than 25 years ago, that *S. pombe* Asp1 shows a strong genetic interaction with components of the Arp2/3 complex but did neither co-sediment nor co-immunoprecipitate with Arp2/3 components [19]. Interestingly, modulation of Arp2/3 components by 1,5-(PP)_2_-InsP_4_ might be evolutionarily conserved. Recently, a proteome-wide quantification of inositol pyrophosphate-human protein interactions recovered all seven mammalian Arp2/3 subunits and components of the WAVE Arp2/3 regulatory complex [52, 64].

How the association of inositol pyrophosphates with Arp2/3 might regulate the complex needs to be analyzed in the future. The complex undergoes a major conformational transition to form a branched actin nucleus, whereas CK666 stabilizes an inactive conformation [48]. Binding of a highly phosphorylated inositol phosphate could therefore influence Arp2/3 conformation, interaction with nucleation-promoting factors, or susceptibility to allosteric inhibition.

### Opposing Roles of Asp1 in Arp2/3-Dependent Actin Regulation in *S. pombe* and *S. japonicus*

Our analysis of actin patches in WT and *asp1Δ S. japonicus* strains demonstrated that SjAsp1 restricts Arp2/3-containing actin patches. Compared with wild-type cells, *Sj asp1Δ* cells exhibited a higher density of cortical actin patches, larger patches, and increased LifeAct fluorescence, whereas patch lifetime remained unchanged. Thus, the increased abundance of actin patches in *Sj asp1Δ* cells is unlikely to result from increased patch persistence. Instead, these findings suggest that patches are initiated more frequently and that greater amounts of F-actin are assembled within each patch.

Regulation of the actin cytoskeleton is important for filamentous morphogenesis in both *S. pombe* and *S. japonicus*. In *S. pombe*, disruption of actin polymerization with latrunculin A blocks invasive mycelial growth [65]. Similarly, the yeast-to-hypha transition in *S. japonicus* is accompanied by extensive reorganization of the actin cytoskeleton, and actin-dependent polarized growth is required for establishment of the highly polarized hyphal growth state [40]. Thus, despite differences in their filamentous growth programs, both fission yeasts require appropriate actin organization to undergo the transition from yeast-like to filamentous growth. Notably, increased Arp2/3-dependent actin assembly similar to that observed in the *Sj asp1Δ* strain has been reported in *S. pombe* when negative regulation of endocytic actin patches is disrupted. Bbc1 and Sla1 proteins constrain Arp2/3-dependent patch assembly by regulating the accumulation of the nucleation-promoting machinery at endocytic sites. In *bbc1Δ sla1Δ* cells, patch-associated LifeAct fluorescence is increased approximately twofold, together with increased recruitment of multiple actin assembly and endocytic regulatory components [66]. This phenotype resembles *Sj asp1Δ*, suggesting that loss of SjAsp1 may similarly release a restraint on Arp2/3-dependent actin assembly.

The altered actin organization in *Sj asp1Δ* may therefore be related to its enhanced invasive growth and hyphal switching. Because actin organization is required for filamentous morphogenesis in both *S. pombe* and *S. japonicus*, increased Arp2/3-dependent actin assembly in *Sj asp1Δ* could create a cytoskeletal state that is more permissive for the transition to polarized hyphal growth. Together, these observations suggest that SjAsp1 normally restricts cortical Arp2/3-dependent actin assembly and that loss of this restraint correlates with an increased propensity to undergo invasive and hyphal growth. Whether enhanced Arp2/3- dependent actin assembly directly drives the increased morphological switching of *Sj asp1Δ* cells remains to be determined.

Our data provide a cellular basis for the species-specific inversion described above and place the difference at the level of Arp2/3 or its immediate regulatory environment. Species-specific networks could determine how engagement by the same inositol pyrophosphate signal is translated into opposite outputs. Together, our results distinguish conservation of signal production, candidate target engagement, and signal interpretation. Asp1 generates a functional inositol pyrophosphate signal in all three fungi, and Arp2/3 emerges as a candidate effector of this pathway. However, the biological meaning assigned to that signal is not fixed: *S. pombe* and *U. maydis* share one regulatory output, whereas *S. japonicus* has reversed it.

## Material and Methods

### 2.1 Strains

Strains used in this study are listed in Table S1. Gene deletion of *Sjasp1^+^* was done via PCR based gene targeting [67]. Briefly, the *Sjasp1Δ* strain was generated via an *S. pombe ura4^+^* cassette [68] with 2 kb long 5’ and 3’ homologue sequences to the 2,8 kb long *Sjasp1^+^* ORF (JaponicusDB, SJAG_00244). The linearized vector was integrated into *S. japonicus* NIG5091 via homologues recombination. Deletion of *Sjasp1^+^* was verified by Southern blot analysis. All plasmids were constructed as previously described [69]. To investigate the actin cytoskeleton in the *S. japonicus asp1* deletion mutant, the *asp1Δ* strain was crossed with the LifeAct strain from the Martin laboratory on SPA medium [70]. The strains were mixed in 15 µl dH₂O and spotted onto the plate. After incubation for 18 h at 25 °C, successful mating was confirmed by light microscopy, and spores were released by β-glucuronidase digestion. The spores were plated onto YE5S medium and subsequently screened by PCR and microscopy to confirm the correct genotypes.

The pJR2-3XL vector containing the thiamine-repressible *nmt1^+^* promoter [71] was used for expression of PPIP5K variants (kinase/phosphatase) from *S. pombe* (SpAsp1), *S. japonicus* (SjAsp1), and *U. maydis* (UmAsp1) in thiabendazole (TBZ; Sigma-Aldrich) assays. Coding sequences were obtained from the respective databases, codon-optimized for expression in *S. pombe* where required, and cloned into the *nmt1^+^*-based expression vector for heterologous expression. Previously described plasmid-expressed *S. pombe asp1^+^* variants were used as controls [23, 24]. *S. japonicus* sequences were retrieved from japonicusdb.org, whereas *U. maydis* sequences were obtained from EnsemblFungi.

### 2.2 Media and growth conditions

If not described differently, *S. pombe* and *S. japonicus* strains were grown at 30 °C on full media (YE5S, 3 % Glucose) or synthetic minimal media (MM, 2 % Glucose) with the appropriate supplements [72]. Cultures were shaken at 150 rpm or grown on 2 % agar plates. *U. maydis* was cultured on plates in complete medium (CM, 1% glucose) or, in the case of liquid cultures, in test tubes on a rotating wheel at 28 °C. Plasmid-borne *Spasp1* and *Sjasp1* variants were expressed via the *nmt1^+^ promoter* [71]. Transformed *S. pombe* cells were grown in MM in the presence of 5 µg/mL (18.8 µM) thiamine (low expression, Sigma Aldrich), in the presence of 0.05 µM thiamine (intermediate expression), or without thiamine (high expression). For serial dilution patch tests either 10^5^ or 10^4^ to 10^1^ logarithmically growing cells were spotted on agar plates and incubated under the conditions described in the figure legends. Invasive growth assays were performed by spotting 10^5^ cells on YE5S followed by incubation (4 days for *S. japonicus* strains, 2-3 weeks for *S. pombe* strains). Adhesive and invasive growth was analyzed as described [24, 73]. To induce mycelial growth in *S. japonicus*, 10^5^ cells were spotted on solid (2% agar) YEMA media [38]. Early mycelial growth was analyzed after 4 days. Full mycelial formation was visible after 7 days of incubation at 30 °C. The Topoisomerase I inhibitor Camptothecin (CPT, Cayman Chemical) induces hyphal development in *S. japonicus* grown in liquid media [74]. Thus, logarithmically growing *S. japonicus* cells were diluted to an OD_600_ of 0.2. CPT was added to a final concentration of 0.2 µM. YEMA und invasive growth plates (YE5S) were scanned with an Epson Perfection V370 Photo scanner. The size of the colonies was determined using the Freehand tool in ImageJ by drawing around them and using the Analyze function. The total number of visible invasive colonies per patch was counted. The Arp2/3 inhibitor CK666 (Sigma-Aldrich) was added to agar plates.

### 2.3 Protein structure prediction and structural comparison

AlphaFold Monomer v2.0 predicted structures were obtained from the Uniprot database (accessed on 26.04.2026). The following UniProt accession numbers were used: *S. pombe* O74429, *S. japonicus* B6JV42 and *U. maydis* A0A0D1BUD3. Pairwise amino acid sequence identities were determined by BLASTP.

### 2.4 [^3^H] Inositol Labeling and HPLC Analysis

Inositol phosphates were determined using [^3^H] Inositol labeling followed by strong anion-exchange (SAX) HPLC analysis as described [35] with little modification. Briefly, *S. japonicus* strains which are inositol auxotroph were grown overnight in liquid PMG medium with appropriate supplements and 10 µM inositol, followed by a dilution to an OD_600_ = 0.05 and grown again overnight in liquid PMG with supplements + 10 µM inositol and + 6 μCi/ml [^3^H]-inositol (Revvity (NET114A005)) at 30 °C. Incorporation of [^3^H]-inositol was allowed for 4-5 cell divisions. *U. maydis* was inoculated into MM medium without inositol (*U. maydis* has an *ino1^+^* gene that encodes inositol synthase and can therefore synthesize inositol de novo,[75]) at 30 °C and 150 rpm and diluted the following day to form a main culture (WT 0.007, *asp1Δ* 0.03), which was labeled with [^3^H]-inositol until the next morning (OD=1). In that case, the incorporation of [^3^H]-inositol was allowed for 5–7 cell divisions. Extraction of inositol polyphosphates was carried out as described [35] and resolved by SAX-HPLC (using a Parti-Sphere SAX 4.6- by 125-mm column; Hichrom). 1 ml fractions were collected for 80 minutes and analyzed by scintillation counting [35]. To quantify the amounts of InsPPs the fractions of each peak were summed. Baseline correction was then performed (in which the mean of the last two measured values, multiplied by the number of fractions of the respective peak, was subtracted from the peak value). The relative ratio of (PP)_2_-InsP_4_/(PP)-InsP_5_ to InsP_6_ was determined and statistically analysed using GraphPad Prism 10.2.3.

### 2.5 Microscopy

Invasive colonies were analyzed by excising agar blocks and placing them on microscope slides. Images were captured using a Zeiss Axiovert 200 fluorescence microscope (Carl Zeiss, Jena, Germany) at 4× magnification. Yeast and hyphal forms of *S. japonicus* were visualized using a Nikon Eclipse Ti confocal fluorescence microscope in bright-field mode at 60× magnification. To visualize the mitochondria, *S. japonicus* cells were stained with 0.1 µg/ml

MitoTracker CMXRos (M7512, Thermo Fisher), in YE5S medium at 30 °C for 30 minutes. Cells were then fixed in 3% formaldehyde at room temperature for 30 minutes followed by microscopy. For *U. maydis*, cells were stained with 15 nM tetramethylrhodamin (TMRE, T669, Thermo Fisher) for 20 min at 28 °C in the dark, then washed once with PBS and plated onto 2% agarose agar pads. To visualize actin in *S. japonicus* WT and *asp1Δ* cells, LifeAct-GFP strains were used. For microscopy of actin in *S. japonicus* and of mitochondria and microtubules in *S. pombe*, *S. japonicus*, and *U. maydis*, a confocal spinning disk microscope (Olympus, Tokyo, Japan) with a 100x oil objective was used. For quantification of actin patch lifetime in *S. japonicus*, the fast scan mode of the Zeiss LSM 880 AiryScan was used with a 63x oil objective.

### 2.6 Programs and Statistical Analysis

To visualize data and to perform statistics GraphPad Prism (version 10.2.3) was used. For generating charts and figures, Biorender, GraphPad Prism and Canvas 14 were used.

Actin patch density, size, fluorescence intensity, lifetime, and spatial distribution were quantified from fluorescence microscopy images of *S. japonicus* WT and *asp1Δ* cells, LifeAct-GFP strains using Fiji/Image. All quantitative analyses were performed using images acquired under identical imaging conditions for the respective comparison. For quantification of actin patch density, the outline of each individual cell was manually delineated as a region of interest (ROI) in Fiji/ImageJ, and the corresponding cell area (µm²) was determined. Actin patches located within the cellular ROI were counted manually. Actin patch density was calculated for each individual cell by dividing the total number of actin patches by the corresponding cell area and is shown as patches/µm². For determination of actin patch size and fluorescence intensity, individual actin patches were manually delineated as ROIs in Fiji/ImageJ. For each patch, the ROI area, mean fluorescence intensity, and integrated density were measured. To correct for local cytoplasmic fluorescence, the mean background fluorescence was measured in a patch-free region adjacent to each individual actin patch. The background-corrected integrated patch fluorescence intensity was calculated as: Corrected integrated density = Integrated density(*patch) − (Area(*patch) × Mean fluorescence(background)). Corrected integrated density values are reported in arbitrary units (a.u.). Patch area is reported in µm². For analysis of actin patch lifetime, actin patches were followed individually via time-lapse fluorescence microscopy recordings from their first appearance to their disappearance. Images were acquired at a temporal resolution of 2.07 s per frame. Patch lifetime (seconds) was calculated from the number of frames during which an individual patch was detectable. Spatial distribution of actin patches was scored by dividing cells were longitudinally into three regions, consisting of two cell end regions and one cell middle region and counting the number actin patches located within the cell middle region.

### 2.7 1,5(PCP)_2_-InsP_4_ pulldown and MS analysis

#### Cell cultivation and lysis

*S. pombe* wild-type cells were grown in PMG medium supplemented with 2% glucose at 30 °C to mid-log phase (OD₆₀₀ = 0.5). A total of 4 L culture (2000 OD units) were harvested (4000 × g, 10 min, 4 °C). Cell disruption was performed adapting a cryogenic cell lysis protocol [76]. In short, Pellets were washed once with 20 mL and twice with 5 mL of ice-cold 50 mM Tris-HCl (pH 7.5), followed by three centrifugation steps (2600 × g, 5 min) to remove as much buffer as possible. Washed pellets were transferred into a 20 mL syringe and extruded into liquid N₂ to generate “yeast noodles” which were stored at −80 °C. Cell disruption was performed using a Retsch MM400 mixer mill. N_2_-cooled 35 mL grinding jars containing a single 10 mm steel ball were loaded with frozen cell noodles (corresponding to 2 L culture per jar). Samples were lysed at 30 Hz for 3 min, repeated six times under continuous cooling. Cell breakage efficiency was verified microscopically.

#### Preparation of biotin-1,5(PCP)_2_-InsP_4_ (b-1,5(PCP)_2_-InsP_4_) matrix and pulldown [77]

Affinity purifications were carried out in a 96-well hydrophilic PVDF Multiscreen plate placed on a Plate Prep 96-well vacuum manifold. Filter plates were pre-washed with 50 µL 70% ethanol followed by two washes with 200 µL PBS. Streptavidin–Sepharose High Performance beads (GE Healthcare, 17-5113-01; binding capacity 5 nmol 1,5(PP)_2_-InsP_4_ per 20 µL beads) were added at 60 µL per well and washed three times with 200 µL PBS. Biotinylated 1,5(PCP)_2_-InsP_4_ (b-1,5(PCP)_2_-InsP_4_) was diluted in PBS to 100 µM and 150 µL of this solution was added to each well and incubated for 20 min at 4 °C and 300 rpm. Controls were processed in parallel by adding 150 µL PBS without 1,5(PCP)_2_-InsP_4_. Beads were washed once with 200 µL PBS and twice with binding buffer (25 mM HEPES pH 7.4, 150 mM NaCl, 0.05% Triton X-100). Meanwhile, the frozen cell powder was resuspended at 1 mg/mL total protein (BCA assay) in binding buffer supplemented with PMSF (final concentration 1 mM). Samples were incubated for 10 min at 4 °C and centrifuged (3000 × g, 10 min, 4 °C). For each pulldown, 150 µL soluble lysate (= 150 µg protein) was added to beads (incubation 1 h at 4 °C, 300 rpm). Beads were subsequently washed six times with 200 µL binding buffer. The filter plate was placed onto a standard 96-well collection plate. Proteins were competitively eluted by adding 50 µL 5mM non-biotinylated 1,5-(PCP)_2_-InsP_4_ in 25 mM HEPES pH 7.4, incubated for 10 min at 4 °C, and centrifuging (500 × g, 1 min). A second elution was performed identically using fresh 1,5(PCP)_2_-InsP_4_ solution and eluates were combined with the first elution. The pulldown was done in parallel using the same concentrations of 3 linked biotinylated 1,5(PCP)_2_-InsP_4_, 1/3-linked biotinylated 5PCP-InsP_5_ (b-5PCP-InsP_5_) and 1/3-linked biotinylated InsP_6_ (b-InsP_6_). In these experiments, proteins were competitively eluted with the corresponding non-biotinylated ligand prior to LC-MS analysis. Generation of the biotinylated and non-biotinylated compounds has been previously described in [52, 77].

##### MS-based quantitative proteomics

MS-based quantitative proteomics was essentially performed as described previously [78] with detailed sample preparation given in [79]. *Sample preparation for LC-MS:* The eluates were supplemented with SDS buffer (final 6% glycerol, 2.4% SDS, 30 mM Tris/HCl pH 7.0), reduced (final 17.8 mM dithiothreitol (DTT), 20 min, 56 °C), alkylated (iodoacetamide (IAA), 8x molar excess to DTT, 15 min, RT, protected from light), and finally underwent on-bead tryptic digestion (100 ng of trypsin in 20 µL 50 mM triethylammonium bicarbonate) after applying a slightly modified sp3 protocol [80] using 100 μg 1:1 mix Sera-Mag SpeedBeads and 50% ethanol (final conc.) for protein precipitation as well as 80% ethanol (3x 200 µL) and acetonitrile (1x 200 µL) as washing solutions. Twenty percent of the peptides were dissolved in 0.1% trifluoracetic acid and subjected to LC-MS analysis. *LC–MS analysis:* For the LC-MS analysis, a Q Exactive Plus Hybrid Quadrupole-Orbitrap mass spectrometer (Thermo Fisher Scientific), coupled with a nano electrospray ionization source connected with an Ultimate 3000 Rapid Separation liquid chromatography (LC) system (Dionex / Thermo Fisher Scientific, Idstein, Germany) equipped with an Acclaim PepMap 100 C18 column (75 μm inner diameter, 25 cm length, 2 mm particle size from Thermo Fisher Scientific) was applied. The mass spectrometer was operated in positive mode and LC was performed using a 120 min gradient (300 nL/min flow rate, gradient from 4 to 40% solvent B, solvent A: 0.1% (v/v) formic acid, solvent B: 0.1% (v/v) formic acid, 84% (v/v) acetonitrile). Capillary temperature and source voltage were set to 250°C and 1.4 kV, respectively. A mass range from 350 to 2000 m/z at a resolution of 140,000 was used for MS survey scans. Automatic gain control (AGC) was set to 3,000,000 and maximum fill time to 80 ms. The ten most intensive peptide ions per survey scan were chosen for high-energy collision dissociation (HCD) fragmentation.

#### Data processing

Raw files were processed with MaxQuant (version 2.2.0.0, Max Planck Institute for Biochemistry, Planegg, Germany) using the *S. pombe* sequence database (UniProtKB, UP000002485, strain972 ATCC24843, downloaded on 11/05/2021, 5151 entries). Carbamidomethylation (Cys) was set as fixed modification, methionine oxidation and N-terminal acetylation as variable modifications. “Match between runs” was enabled and peptide/protein false discovery rate (FDR) was 1%.

Statistical analysis was performed based on experiment-pairwise median log_2_(fold change) normalized MaxQuant protein group intensities using the “R” (v4.2.2) programming language after removing potential contaminants, reverse hits, and proteins only identified by modified peptides. Testing for significance, by applying a 5% permutation-based FDR cutoff in the differential analysis (e.g., 1,5(PCP)_2_-InsP_4_ vs. control), was performed using the “Significance Analysis of Microarrays” (SAM) analysis method [81] within the Siggenes package. For this approach, a minimum of three valid values had to be present in at least one group (1,5(PCP)_2_-InsP_4_ or control), data were log2 transformed to reach a normal distribution like data structure, and missing values were filled in with random values from sample wise downshifted normal distributions (0.3 s.d. width, 1.8 s.d. downshift).

### Structural visualization

The structure of the *S. pombe* Arp2/3 complex (PDB ID: 6W17) was visualized using UCSF Chimera version 1.18 [57]. The bound nucleation-promoting factor and actin molecules were removed for clarity, and individual Arp2/3 subunits were colour-coded for visualization

## Author Contributions

Conceptualization, U.F.; Methodology, E.K., A.A.-R, L.J., A.S., V.E., S.M.B., L.vW., J.P., and T.L.; Software, E.K., L.J., T.L. and I.S.; Validation, E.K., A.S., S.M.B., A.A.-R., V.E., and L.J.; Formal Analysis, E.K., A.S., L.J., A.A.-R., T.L., V.E., and I.S.; Investigation, E.K., A.S., A.A.-R., L.J., L.vW., V.E., and J.P.; Resources, U.F., D.F., K.S., and M.F.; Writing-Original Draft Preparation, U.F.; Writing-Review and Editing, U.F., L.J., A.S., L.vW., and E.K.; Visualization, E.K., L.J., T.L., and I.S.; Supervision, U.F., D.F., and M.F.; Project Administration, U.F.; Funding Acquisition, U.F., D.F., M.F., and A.S.

## Acknowledgements

We are grateful to the Center for Advanced Imaging (CAi) at Heinrich Heine University Düsseldorf for their support with microscopy. We thank S. Bollé (HHU) for assistance with Figure 4, and L. Geerkens (HHU) for the serial dilution patch test in Sup Fig. 3A. We are also grateful to J. Hegemann (HHU) for carefully reading the manuscript. We thank Kathleen Gould (Vanderbilt University, USA), Sophie Martin (University of Geneva, Switzerland), Phong Tran (Institut Curie, France), and Hironori Niki (National Institute of Genetics, Japan) for providing strains. The funders had no role in study design, data collection and analysis, decision to publish or preparation of the manuscript.

