## Supplementary Figures for "Species Context Reverses PPIP5K Control of Fungal Morphogenesis and Actin Organization"

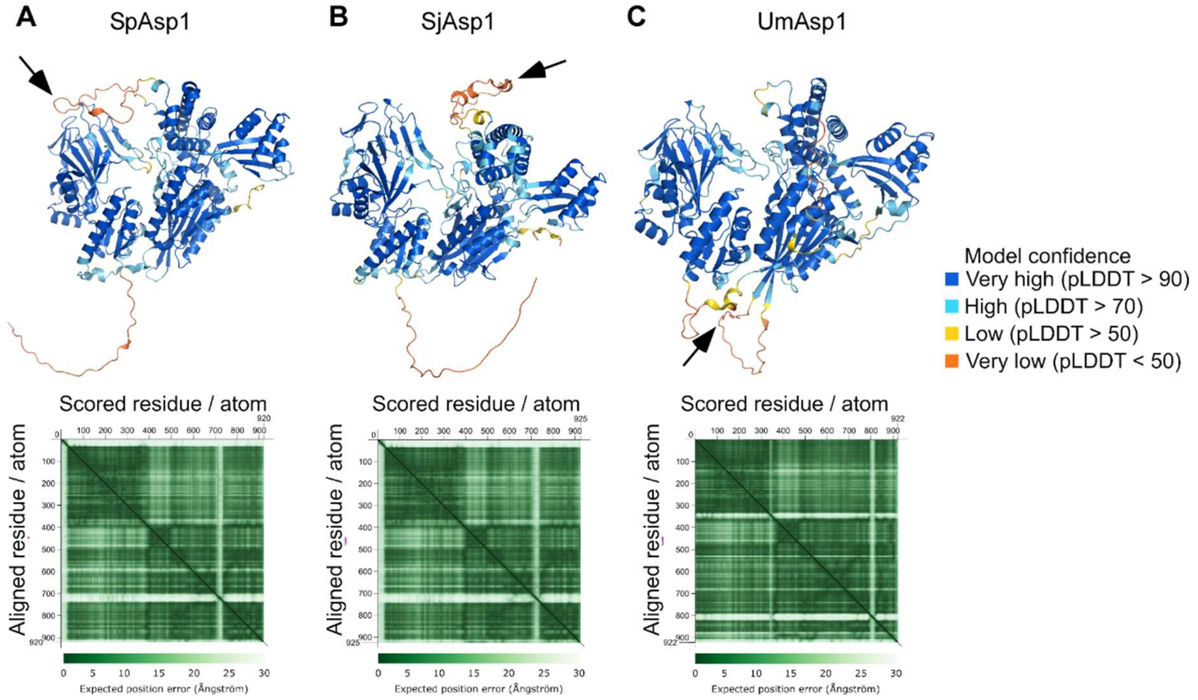


**Supplementary Fig. S1. Confidence assessment of AlphaFold2-predicted structures of PPIP5K homologs (A-C)** AF2-predicted structures of **(A)** SpAsp1, **(B)** SjAsp1, and **(C)** UmAsp1 coloured according to the predicted Local Distance Difference Test (pLDDT), with blue indicating very high confidence (pLDDT > 90), cyan high confidence (pLDDT > 70), yellow low confidence (pLDDT > 50), and orange very low confidence (pLDDT < 50). Black arrows indicate the flexible regions within the phosphatase domain that are predicted with low local confidence, corresponding to the region around residue 700 in SpAsp1 and SjAsp1 and the region around residue 800 UmAsp1. The lower panels show the Predicted Aligned Error (PAE) matrices for each model. PAE is a measure of the confidence in the relative position of two residues within the predicted structure, providing insight into the reliability of relative position and orientations of different domains.

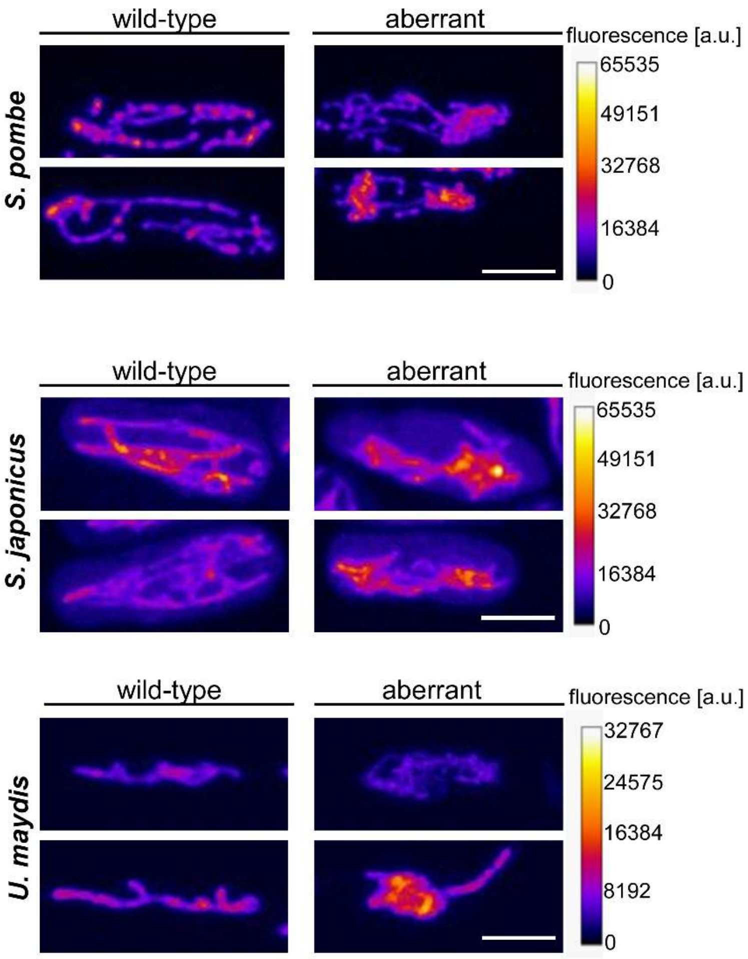


**Supplementary Fig. 2 Species-specific effects of 1,5(PP)$_2$-InsP$_4$ on mitochondrial organization.** Representative images illustrating the "wild-type-like" and "aberrant" mitochondrial morphology categories in *S. pombe* (top panel), *S. japonicus* (middle panel), and *U. maydis* (bottom panel). Wild-type-like mitochondrial phenotypes are characterized by long, tubular mitochondria extending along the longitudinal axis of the cell, whereas aberrant mitochondria exhibit reduced tubularity, shorter structures, or mitochondrial aggregates. Images were displayed using the "Fire" lookup table (LUT) in ImageJ, which maps fluorescence intensity values to a pseudo color scale, with dark blue/purple representing lower and orange/light yellow representing higher fluorescence intensities. Scale bars: 5 µm

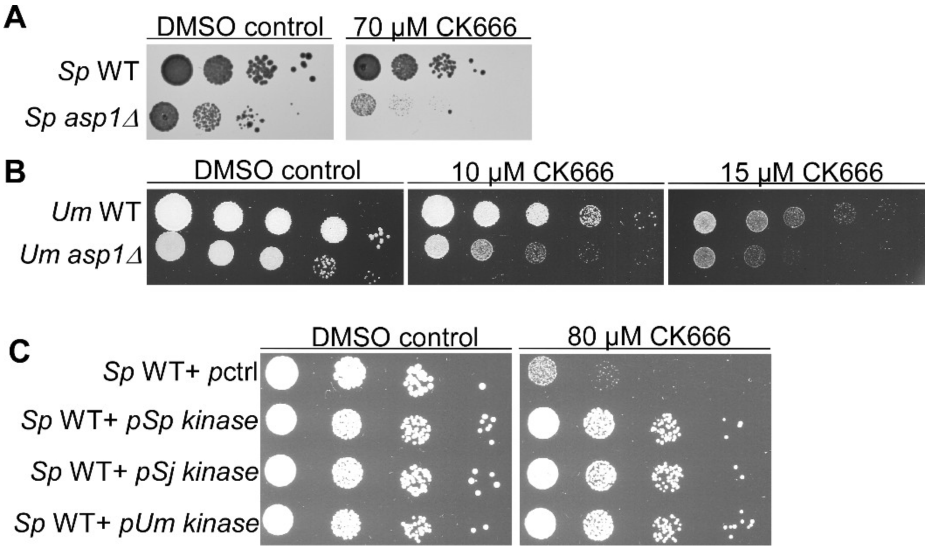

**Supplementary Fig. 3. Asp1 deficiency increases sensitivity to the Arp2/3 inhibitor CK666 in *S. pombe* and *U. maydis*.** (**A**) Serial dilution patch test assay ($10^5$–$10^1$ cells) of *Sp* WT and *Sp asp1Δ* strains on YE5S medium supplemented with DMSO (DMSO control) or the Arp2/3 inhibitor CK666. Plates were incubated at 25 °C for 3 days. (**B**) Serial dilution patch test assay ($10^5$–$10^1$ cells) of *Um* WT and *asp1Δ* strains on CM medium containing DMSO (DMSO control) or CK666. Plates were incubated at 28 °C for 3 days. (**C**) Serial dilution patch test ($10^4$–$10^1$ cells) of *Sp* WT transformants carrying a control plasmid (p ctrl), or plasmids with the *S. pombe* kinase domain (p*Sp* kinase, amino acids 1-364), the *S. japonicus* kinase domain (p*Sj* kinase, amino acids 3-366) or the *U. maydis* kinase domain (p*Um* kinase, amino acids 1-336) on plasmid-selective and promoter-derepressing MM medium supplemented with either DMSO (control) or CK666 (80 µM). Plates were incubated at 25 °C for 6 days.

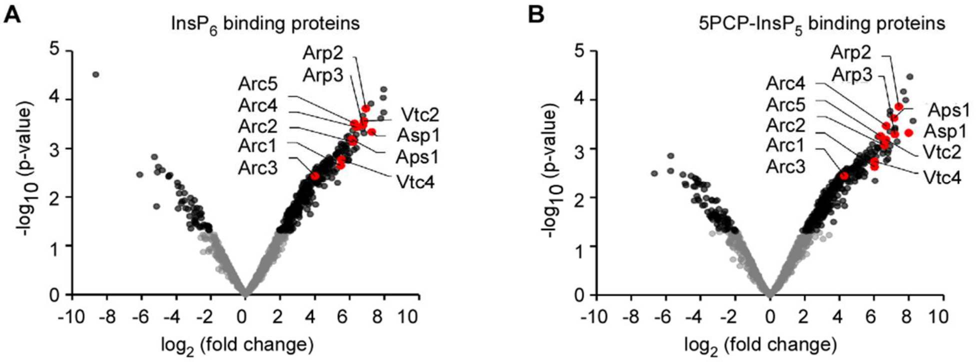


**Supplementary Figure S4. Identification of $InsP_6$- and 5PCP-$InsP_5$ binding proteins in *S. pombe*. (A, B)** Volcano plots showing proteins enriched on b-$InsP_6$- (A) or b-5PCP-$InsP_5$-coated beads (B) relative to control beads. Bound fractions were washed and eluted with an excess of non-biotinylated $InsP_6$ or non-biotinylated 5PCP-$InsP_5$, respectively, and identified by quantitative LC-MS-based proteomics. Statistical analysis was performed as described for Fig. 8D. Proteins of interest are highlighted in red and labelled.

### Table S1: List of strains

*S. pombe* strains

| name | genotype | source |
|---|---|---|
| #605 | h-, his3-D1, ade6-M210, leu2-1, ura4-D18 | K. Gould |
| #1511 | h+, asp1$^{D333A}$::kanR, his3-D1, ade6-M210, leu2-1, ura4-D18 | U. Fleig |
| #1579 | h+, asp1$^{H397A}$::kanR, his3-D1, ade6-M210, leu2-1, ura4-D18 | U. Fleig |
| #2908 | h-, cox4-RFP:LEU2 ade6-m210 leu1-32 ura4-D18 | P. Tran |
| #3327 | h- ade6+:patb2:sfGFP-atb2::hphMX | S. Martin |
| #3601 | h-,leu1-32::cox4-mRFP::LEU2 ade6+:patb2:sfGFP-atb2::hphMX his3-D1 ura4-D18 | U. Fleig |
| #3627 | h-,leu1-32::cox4-mRFP::LEU2 ade6+:patb2:sfGFP-atb2::hphMX asp1D333A::kanR | U. Fleig |

*S. japonicus* strains

| name | genotype | source |
|---|---|---|
| #2100 | h+, ura4$^{Sj}$-D3 | H. Niki |
| #2466 | h+, asp1Δ::ura4$^{+}$, ura4$^{Sj}$-D3 | U. Fleig |

| | | |
|---|---|---|
| #3801 | h-, GFP-atb2::ura4$^{+}$, ade6$^{sj}$-domE | S. Martin |
| #3803 | h-, LifeAct-GFP::ura4$^{+}$, mCherry-atb2::ura4$^{+}$, ade6$^{sj}$-domE | S. Martin |
| #3828 | h+, asp1Δ::ura4$^{+}$, GFP-atb2::ura4$^{+}$ | U. Fleig |
| #3851 | LifeAct-GFP::ura4+, asp1Δ::ura4$^{+}$ | U. Fleig |

*U. maydis* strains

| **name** | **genotype** | **Source** |
|---|---|---|
| AB33 | a2, $P_{nar}$:bW2bE1 | M. Feldbrügge |
| Uma958 | a2 $P_{nar}$:bW2bE1 asp1Δ::HygR | M. Feldbrügge |
| Uma498 | AB33_otef-Tub1-gfp-cbx | M. Feldbrügge |
| Uma991 | AB33_otef_GTub1/asp1D | M. Feldbrügge |
